# RUNX2 promotes epigenetic WNT signaling in inflamed intestinal epithelial cells

**DOI:** 10.1101/2025.11.17.688458

**Authors:** Rodolfo I Cabrera-Silva, Zachary S. Wilson, Jael Miranda, Shuling Fan, Michael Dame, Shrinivas Bishu, Jason R Spence, Jennifer C Brazil, Justin Colacino, Asma Nusrat, Charles A Parkos

## Abstract

Ulcerative colitis (UC) is characterized by chronic mucosal inflammation, recurrent epithelial injury, and impaired colonic mucosal wound healing. While WNT/β-catenin dysregulation has been reported in UC, the mechanisms of such abnormalities remain unclear. To investigate epithelial intrinsic alterations associated with UC, we performed single-nucleus RNA-seq (snRNA-seq) and ATAC-seq (snATAC-seq) multiomics on human primary colonic epithelial cells (colonoids) from healthy donors and patients with inactive or active UC. Colonoids were cultured in a 3D matrix recapitulating crypt base cells or grown as 2D monolayers in differentiation medium to recapitulate luminal epithelial cells. Colonoids from active UC had a unique cell population with elevated CTNNB1 and reduced APC expression. Chromatin profiling identified enrichment of RUNX2 motifs in this UC-associated cell population. Active UC colonoids exhibited reduced OLFM4 expression in 3D and the differentiation marker VIL1 in 2D, suggesting impaired self-renewal and maturation. RUNX2 inhibition using CADD522 reduced β-catenin levels in 3D colonoids and restored VIL1 expression and junctional β-catenin localization in 2D cultures. These findings reveal an intrinsic defect in epithelial renewal in UC, driven in part by RUNX2-dependent WNT dysregulation. Our study identifies RUNX2 as a transcriptional regulator of epithelial stem cell function and WNT signaling in the inflamed human colon.

**Graphical Abstract summary:** Single-nucleus RNA and ATAC sequencing of UC patient-derived colonoids reveals a RUNX2-associated WNT signature in active inflammation. Elevated β-catenin and reduced *OLFM4* and *VIL1* expression indicate impaired self-renewal and differentiation. Pharmacologic inhibition of RUNX2 restores epithelial maturation, identifying RUNX2 as a key regulator of epithelial dysfunction in UC.

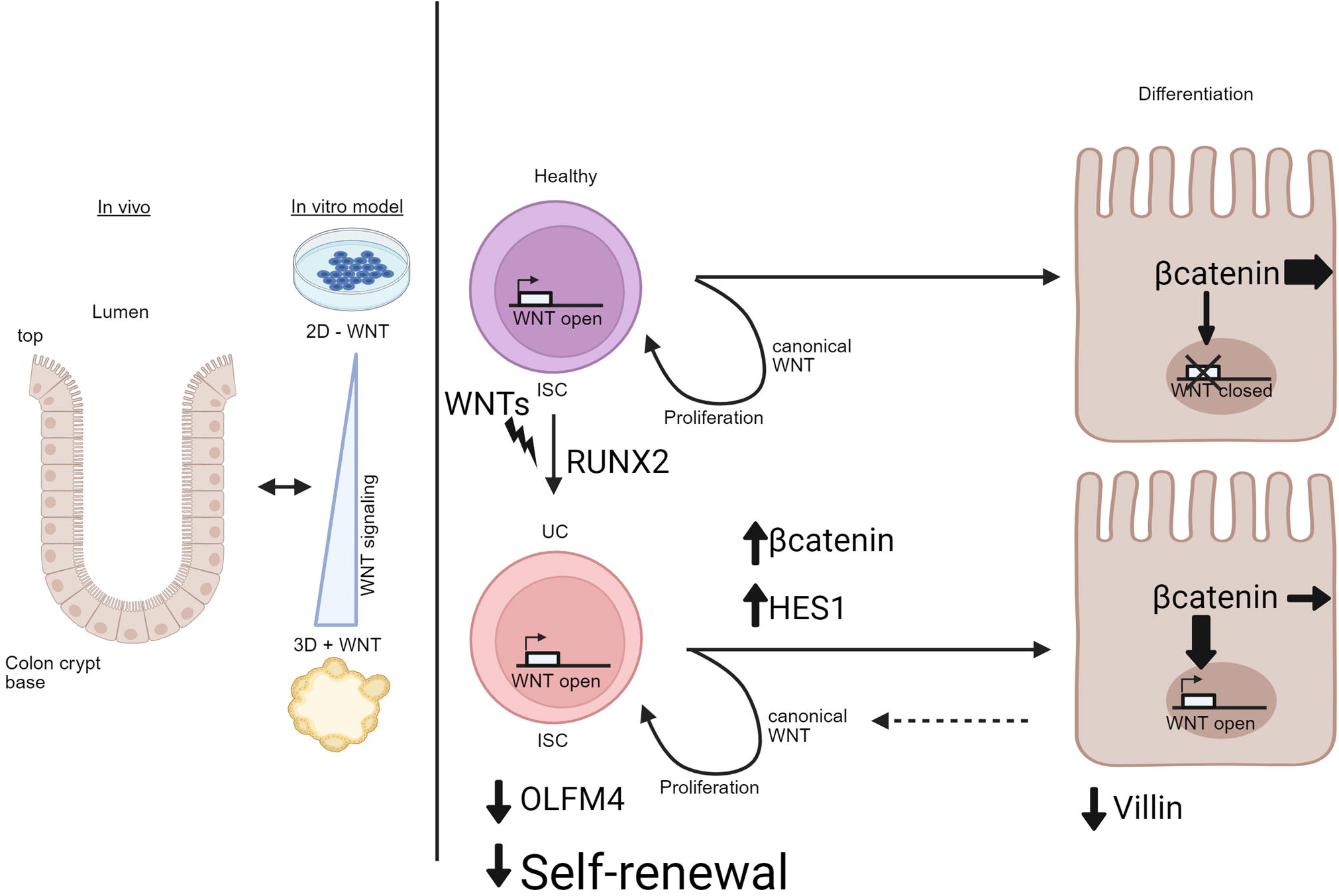

## Introduction

Intestinal epithelial cells (IECs) function as a critical barrier in the gut, restricting entry of potentially harmful substances and microbes while allowing selective passage of necessary ions and nutrients (1). The colonic epithelium is organized into invaginated structures referred to as crypts, which are spatially and functionally compartmentalized into crypt base and crypt top. Cells at the crypt base include highly proliferative cells that replenish the epithelium ∼ 3-5 days (1). In contrast, cells at the crypt top are terminally differentiated and ultimately undergo programed cell death to maintain homeostatic turnover. These distinct compartments support specialized functions, including regulation of ion and water transport, extracellular matrix remodeling, and crosstalk with immune and stromal cells in the lamina propria. Disruption of tightly regulated epithelial renewal and differentiation processes compromises epithelial barrier and impairs mucosal wound repair, both of which are implicated in the pathogenesis of Inflammatory Bowel Disease (IBD).

IBD, encompassing Crohn’s disease (CD) and Ulcerative colitis (UC), is a chronic inflammatory disorder of the gastrointestinal tract characterized by relapsing mucosal injury, which affects 1.6 million people in the United States with an estimated ∼ 70,000 new cases diagnosed annually (2, 3). Elevated epithelial β-catenin activity has been reported in inflamed IBD mucosal tissues (4). Under homeostatic conditions, WNT signaling plays a central role in regulating intestinal epithelial development and self-renewal (5). However, during chronic inflammation, immune cells can secrete soluble WNT ligands, leading to aberrant β-catenin activity and dysregulated epithelial proliferation/differentiation (6–9). Thus, chronic inflammation perturbs critical self-renewal processes in crypt base epithelial stem cells (10). The importance of WNT/β-catenin signaling in intestinal regeneration is highlighted by a recent study demonstrating that FZD5, a component of WNT/β-catenin signaling, regulates epigenetic signaling as well as self-renewal and plasticity in intestinal epithelial cells (11).

In addition to changes in epithelial renewal and proliferation, dysregulation of chromatin accessibility has more recently been linked with disease pathogenesis in UC (12, 13). Chromatin, consisting of DNA wrapped around histones, can adopt “open” or “closed” conformations depending on various factors, including DNA methylation and histone modifications, including acetylation, methylation, or phosphorylation. Transcription factor binding, which plays a vital role in gene regulation, is therefore dependent on both chromatin structure and DNA accessibility. Recent studies suggest that epigenetic modifications of DNA and histones are critical regulators of stem cell self-renewal and colonic epithelial integrity. In UC, disruption of these tightly controlled mechanisms may impair the regenerative capacity of epithelial cells and compromise mucosal homeostasis (13). To better understand how such epigenetic and transcriptional changes contribute to disease, multiomics approaches combining single nuclei RNA sequencing (snRNA-seq) with assay for transposase-accessible chromatin sequencing (snATAC-seq), have emerged as powerful tools for linking transcription factor activity with gene regulatory dynamics and altered cell phenotypes across tissues.

In this study, we performed snRNA-seq and snATAC-seq on primary human colonic epithelial organoids (colonoids) to uncover epigenetic and transcriptional mechanisms contributing to epithelial homeostasis and dysfunction in UC. We identified an active UC-specific colonic epithelial population characterized by elevated *CTNNNB1* (β-catenin) expression, reduced *APC*, and increased chromatin accessibility at RUNX2 binding motifs. These cells also exhibited impaired differentiation, evidenced by reduced expression of *VIL1* (Villin) and increased β-catenin levels. Notably, treatment with the RUNX2 inhibitor CADD522 restored epithelial differentiation markers and decreased β-catenin levels, supporting a functional role for RUNX2 in driving WNT-mediated self-renewal defects. Together, our findings reveal an epigenetically driven WNT/RUNX2 signaling axis that disrupts epithelial renewal in active UC. Better understanding of how dysregulated epithelial crypt-luminal axis differentiation and impaired barrier function can be targeted/reversed could provide novel therapeutic targets for resolving chronic inflammation in individuals with UC.

## Results

### In vitro differentiation of human colonoids

UC is characterized by chronic colonic mucosal inflammation and epithelial injury, which is associated with defective epithelial repair that further contributes to compromised epithelial barrier function. To understand the epithelial dysregulation in UC, we modeled crypt to luminal differentiation using colonoids derived from healthy and active or inactive UC donors (**Table 1A**). To model proliferative (undifferentiated) colonic epithelium, epithelial cells isolated from colonic biopsies of healthy and UC donors were cultured in Matrigel with WNT-rich medium to generate 3D colonoids enriched in intestinal stem cells (ISCs) characteristics/crypt base phenotype. Additionally, to model differentiated, luminal-like epithelial cells, biopsy-derived epithelial cells were cultured as polarized monolayers in WNT-depleted conditions (2D colonoids) (14). 3D colonoids grow as spherical undifferentiated epithelial cell aggregates, as shown by ubiquitous expression of *LGR5* and *OLFM4* and limited expression of mature epithelial differentiation markers (**Figure 1A, Figure S1**). In contrast, 2D colonoids form a single layer epithelial sheet with many differentiated enterocytes expressing Villin *(VIL1*) but not LGR5 (**Figure 1B, Figure S1**). Interestingly, *OLFM4*/*BMI1* expression was elevated in 2D differentiated colonoids, likely due to a population of +4 progenitors filling the ISC niche as has been described for other models (**Figure 1B, Figure S1**) (15–20).

**Figure 1:**
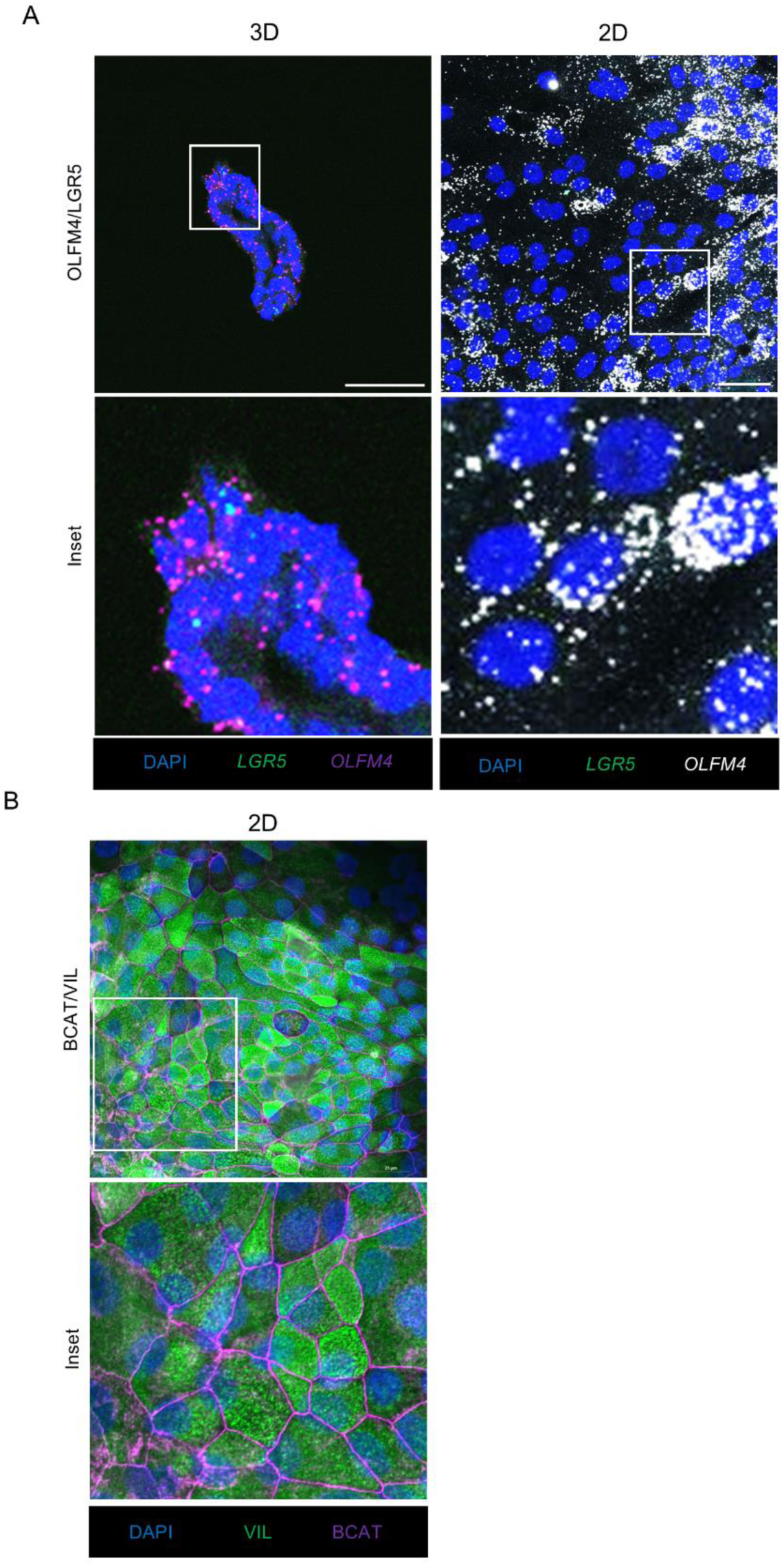
Human colonoids in 3D and 2D conditions. (A) In situ hybridization (RNAscope) of OLFM4/LGR5 mRNA for human colonoids in 3D and 2D conditions (representative of N=3 independent experiments). (B) Immunofluorescence for indicated markers in human colonoids (representative of N=3 independent experiments).

### snRNA-seq identification of discrete epithelial cells clusters in 3D and 2D colonoids

To characterize changes in the intestinal epithelium derived from UC donors compared to healthy donors we performed Multiomic sequencing, joint ATAC-seq and RNA-seq, for 36,777 cells obtained from ‘2D’ and ‘3D’ colonoids (**Figure 2A**). The experimental cohort included two healthy individuals (Healthy 1 and Healthy 2), one UC donor with active disease (active UC), and one UC donor without active disease (inactive UC) (**Table 1A**). Principal component analysis (PCA) from RNA sequencing revealed that cells from the active UC donor colonoids in the 3D condition were distinct compared to healthy and inactive UC samples suggesting fundamental differences in the transcriptional profile in the IECs obtained from actively inflamed mucosa (**Figure 2B, upper panel**). In contrast, this change in transcriptional profile is not observed in 2D colonoids (**Figure 2B, lower panel**), highlighting inflammation-driven changes in the proliferative/stem populations. Following data integration, we identified 11 clusters indicating varying states of epithelial differentiation (**Figure 2D, Figure S2**). As expected IECs cultured as polarized 2D monolayers (differentiated cells) expressed elevated crypt top markers (*REG4, GUCA2A, KRT18, GPA33, CDH17*) while proliferative/stem cells cultured in 3D conditions expressed elevated stem cell markers (*LGR5, LRIG1, SMOC2, EPHB2*) (**Figure 2E-F**). Importantly, in 3D colonoids we identified a cluster exclusive to the active UC donor (‘UC-specific’), along with a cluster of relatively undifferentiated epithelial cells (‘Undifferentiated’) in the 2D condition that might represent a population of quiescent epithelial cells (**Figure 2C-D**). Cells from other epithelial lineages were not found based on current markers including enteroendocrine and BEST4+. In examining various differential gene expression profiles, we found a surprising cluster containing microfold cell (M-like) specific markers (*CCL20, TNFAIP2, ANXA5, MARCKSL1*) which is present in 2D colonoids regardless of disease status/origin (**Figure S3B**). M cells (microfold cells) are predominantly located in the follicle associated epithelial cells overlying lymphoid follicles in the distal small intestine. Sparse M-like cells have been identified over isolated lymphoid aggregates in the colon. These cell types are not observed in vitro in the absence of specific factors to induce M cell phenotypes (21, 22). However, we identified an M-like cluster that abundantly expresses chemoattractants including *IL8, CCL20, and CCL15* as well as fetal regenerative genes such as *TNFRSF12A, LAMC2, CD44, and LY6D* in 2D cultured colonoid monolayers (**Figure S3A & C**). Interestingly, the UC-specific cluster displayed high expression of fetal regenerative genes *YAP1* and *ANXA5* (**Figure S3C**). Altogether, data highlight the presence of a unique cluster identified in inflamed UC 3D colonoids.

**Figure 2:**
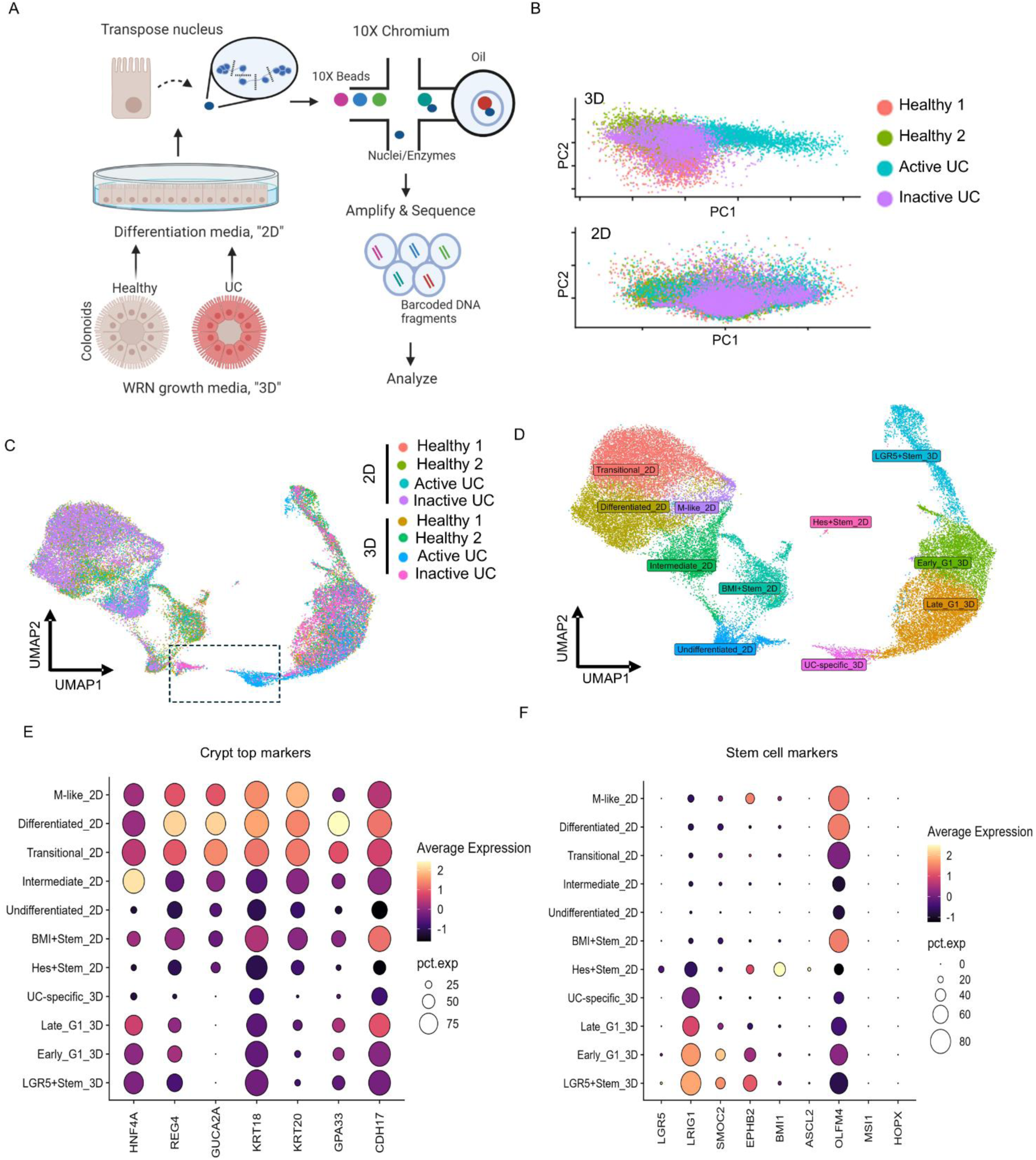
RNA sequencing of colonoids in 2D and 3D from UC individuals. (A) Schematic for multiome (ATAC + GEX) analysis on human colonoids. (B) PCA analysis of 3D and 2D samples. (C-D) UMAP analysis of 3D and 2D samples by (C) sample and (D) cell clusters. (E) Dotplot analysis of top 10 genes in each cell cluster. (F-G) Dotplot analysis of (F) select crypt top markers and (G) select stem cell markers in each cell cluster.

To validate our 3D and 2D designations and to determine how cell populations were related to each other, trajectory analysis was used to identify the differentiation potential of each cell subset in our single-cell dataset. Pseudotime analysis unbiasedly confirmed 3D colonoids transitioning into 2D differentiated populations. Furthermore, we were able to identify ‘LGR5+ Stem’ in 3D that were starting points of differentiation, and ‘M-like’, ‘Undifferentiated’, and ‘Differentiated’ clusters in 2D as terminal points of differentiation (**Figure S4A-C**). Gene expression trends confirmed the increase of mature and M-like cell markers and the decrease of stem cell markers with increasing pseudotime (**Figure S4D**). Pseudotime analyses demonstrated that cells from the ‘UC-specific’ cluster preferentially progresses toward the ‘Undifferentiated’ terminal endpoint (**Figure S4C**). These findings suggest a unique fate for specific epithelial cells derived from active UC intestinal tissues that differs from the otherwise normal differentiation trajectory. Additionally, pseudobulk Gene Set Enrichment Analysis (GSEA) was performed on 2D vs 3D for healthy or active UC samples to identify changes in differentiation (**Figure S5**). For IECs from inflamed UC mucosa, the most upregulated pathway was epithelial-mesenchymal transition (EMT). In contrast, this pathway was ranked 9th for the healthy samples. In addition, p53 and TGF-β pathways were only observed for epithelial cells from the healthy samples. This analysis suggests important alterations in the active UC IECs that impact transcriptional activity (**Figure S5**).

### snATAC seq of primary human colonoids to determine regions of open chromatin

To determine epigenetic changes during differentiation from the ‘3D colonoid’ to the ‘2D colonoid’ state, we analyzed regions of open chromatin (euchromatin) using snATAC-sequencing. Using the ArchR package, we analyzed all samples individually for gene scores based on 500 bp gene tiles. We generated an integrated model for all samples using ArchR’s iterative latent semantic index (LSI) dimensional reduction and Harmony to identify selectively open regions at a single cell resolution among all samples (23) (**Figure 3A-B**). In examining the number of open region peaks per cluster, we report that peak calling of gene tiles estimates fewer peaks than expected for ‘LGR5+ Stem’, ‘HES+ Stem’, and ‘BMI+ Stem’ clusters (**Figure 3C**). This was unexpected based on a previous study reporting that ISCs are expected to have far more open chromatin (24). To resolve this, we increased the false discovery rate (FDR) threshold for the peak calling function and found that at high FDR the clusters with the most peaks were ‘LGR5+ Stem’ and ‘HES+ Stem’ (**Figure 3D**). This would suggest that while more genomic regions are open in these populations, they are stochastic and not selectively open regions. Additionally, these findings support our snRNA-seq analyses in that there was a similar ‘UC-specific’ population identified in the active UC 3D sample for snATAC-seq (**Figure 3B**).

**Figure 3:**
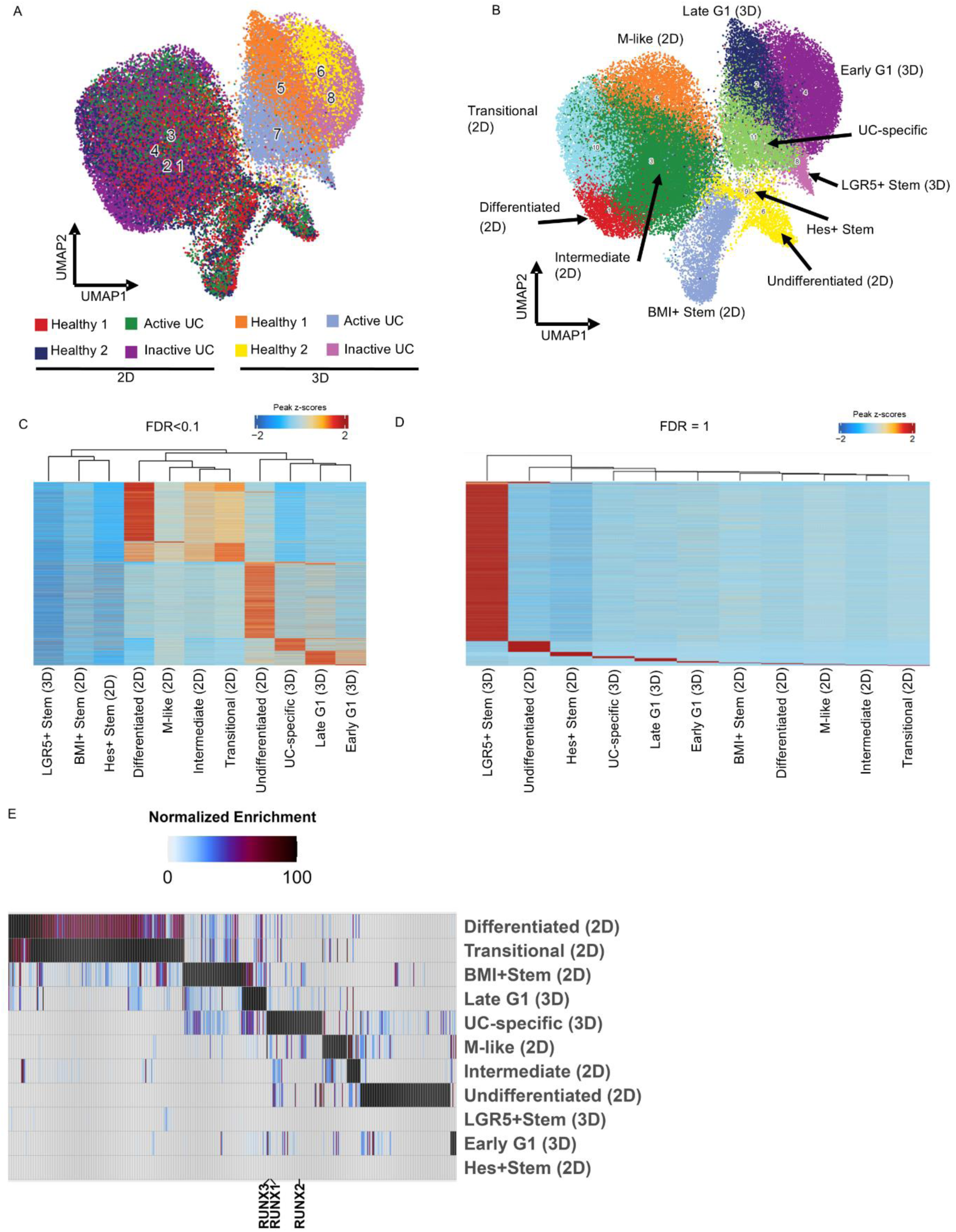
ATAC Sequencing of colonoids in 2D and 3D from UC individuals. (A-B) UMAP analysis of 3D and 2D samples by (A) sample and (B) cell clusters using ArchR pipeline. (C-D) Heatmap analysis of gene peaks per cell cluster based on false discovery rate (FDR) of (C) less than 0.1 and (D) 1. (E) Motif analysis based on cell cluster with indicated motifs above.

To verify that identified populations were similar to clusters found by snRNA sequencing, we looked at integrated gene activity, which combines peak activity with gene expression (**Figure S6-7**). We demonstrate that 2D colonoids have similar differentiated populations as typified by ‘crypt top’ markers (*KRT20, CDH17, GPA33*) and by M-like cell marker *TNFAIP2* (**Figure S6A-B**). Interestingly, in the ‘LGR5⁺ Stem’ and ‘BMI1⁺ Stem’ populations, we detected gene activity associated with enteroendocrine and BEST4⁺ cell markers, lineages that were not observed in the snRNA-seq dataset (Figure S6A). This suggests that these stem cells are epigenetically primed to differentiate into specific lineages through chromatin accessibility but fail to do so under the current in vitro conditions. Pseudotime trajectory analysis of gene activity and integrated gene activity (combined RNA and ATAC seq) shows a progressive decrease in *LGR5* along with increased *VIL1*/*MUC2* activity in 2D differentiated epithelial cell populations (**Figure S7**). A concomitant, although smaller, trajectory of gene activity for *VIL1*/MUC2 was observed moving from 3D ‘LGR5+ Stem to 3D ‘Early G1, ‘Late G1, and ‘UC-specific’ cells. Interestingly, *OLFM4* gene activity was elevated in 2D ‘BMI+ Stem’ and integrated gene activity was found to be elevated in multiple spots of the trajectory pseudotime which may indicate that the chromatin for *OLFM4* is more frequently accessible during epithelial differentiation. Unlike *OFLM4*, *LGR5* gene activity was elevated early in the 3D ‘LGR5 Stem’ cells suggesting that *LGR5* activity is promoted at specific time points in ISCs.

### WNT signaling is enhanced in UC colonoids

After obtaining a set of peaks that uniquely identify ‘active UC-specific’ cells, enriched motifs belonging to various transcription factors were obtained for each cell cluster (**Figure 3E**). The active UC-specific cluster was found to be particularly enriched for motifs corresponding to RUNX family of transcription factors (RUNX1-3), which are WNT-responsive transcription factors that also contribute to WNT signaling (25). We focused on RUNX2 which is not well-characterized in IECs but is important for osteoclast differentiation and cancer development (25). Furthermore, we looked at a larger database of inflamed epithelial cells from UC individuals to determine if there are changes in RUNX transcription factors or WNT signaling in active UC colonoids compared to inactive UC colonoids (26). We confirmed that in discrete populations of inflamed and noninflamed epithelium from UC individuals there is increased expression of *CTNNB1*, a downstream component of WNT signaling, along with increased gene expression of its regulatory protein *APC* and *RUNX1-3* (**Figure S8-9**). Interestingly, while motifs for all RUNX family members are enriched in the UC-specific cluster, when performing Differentially Expressed Genes (DEG) analysis, only *RUNX2* mRNA was upregulated (log2 Fold Change > 1.5) in the active UC sample and therefore is the focus of our subsequent analysis (**Figure S10**). Based on a WNT-responsive transcription factor being upregulated in ‘UC-specific’ cells and these cells being only found in colonoids cultured in conditioned media containing WNTs, we focused our investigation on downstream WNT signaling within this cluster and the overall WNT signaling in the active UC colonoids.

Canonical WNT signaling is initiated by WNT ligands binding to Frizzled (FZD) receptors and co-receptors LRP5/6, leading to activation of Disheveled (DVL) proteins and stabilization of β-catenin, which translocates to the nucleus and interacts with TCF/LEF transcription factors to regulate target gene expression (27). WNT signaling is counter-regulated by a destruction complex composed of proteins such as AXIN and APC, which promote β-catenin phosphorylation and subsequent proteasomal degradation, thereby preventing its nuclear accumulation and transcriptional activity (27). Consistent with previous reports of elevated WNT/β-catenin signaling in individuals with UC (4, 28), we observed marked upregulation of β-catenin mRNA (*CTNNB1*) and a concomitant downregulation of *APC* mRNA within the active UC-specific cluster. These changes were further confirmed by pseudobulk analysis of active UC samples under both 2D and 3D culture conditions (**Figure 4A-D**). A shift in *DVL1* & *DVL3* was noted with *DVL2* being upregulated in the active UC-specific cluster, along with a concomitant downregulation of LRP and FZD proteins (**Figure 4C**). Interestingly, analysis of single nucleotide polymorphisms (SNPs) within the *CTNNB1* transcript revealed a greater number of SNPs in both active and inactive UC-derived colonoids, based on comparison with two comprehensive SNP databases covering the *CTNNB1* genomic region (29, 30) (**Figure 5A-B**). Importantly, polygenic risk based on the combined effect of multiple common polymorphisms provides a more accurate indicator of susceptibility to complex diseases such as UC than single monogenic variant (31). Additionally, we observed a higher number of SNPs in active UC colonoids cultured under 2D conditions compared to 3D, suggesting increased genomic variation in more differentiated epithelial cells (**Figure 5A-B**). Altogether, these data suggest that active UC is associated with epigenetic alterations that modulate colonic WNT signaling in a RUNX2 dependent manner.

**Figure 4:**
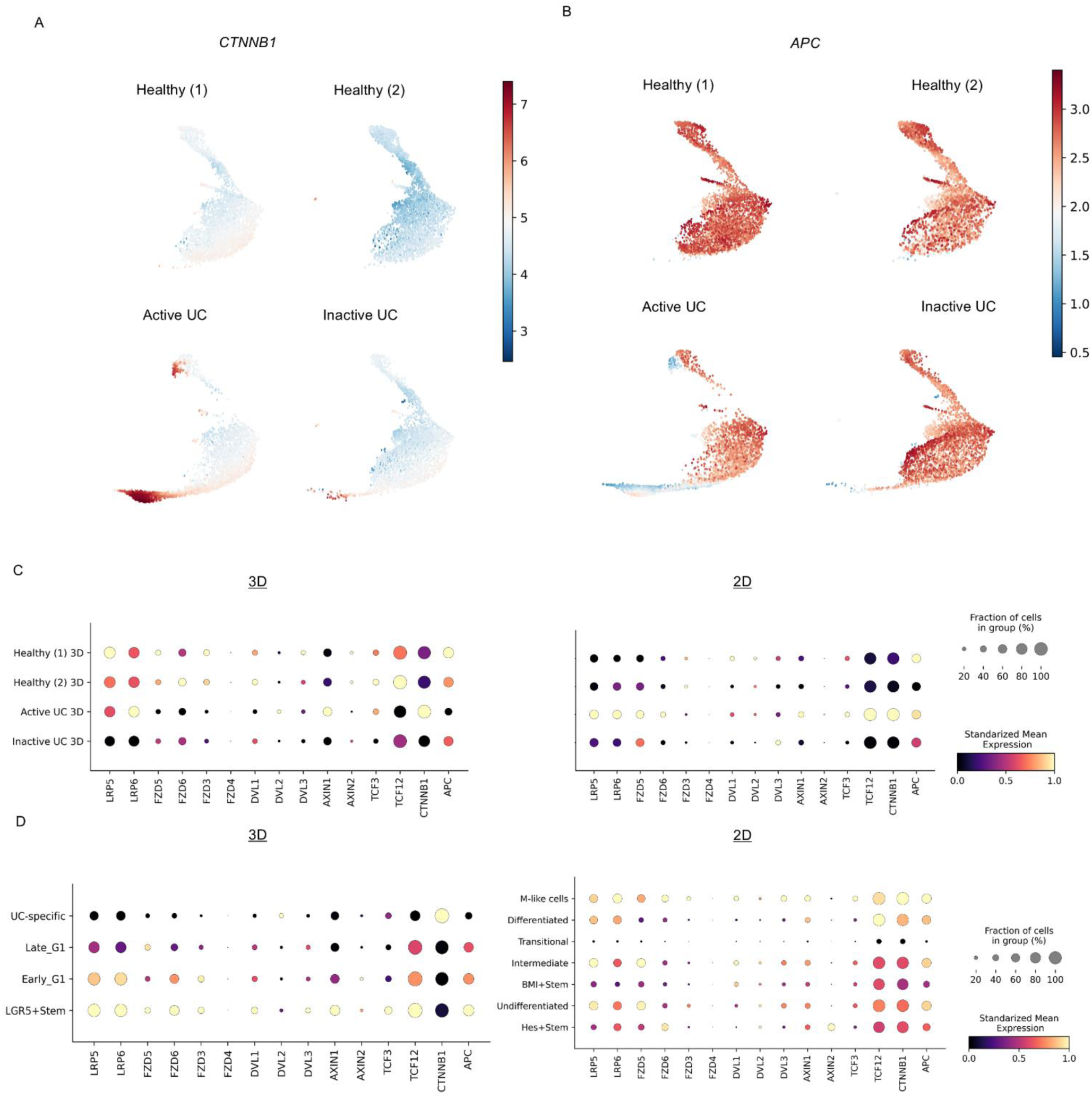
WNT signaling in UC colonoids is upregulated. (A-B) UMAP analysis of gene expression for (A) CTNNB1 and (B) APC in 3D samples. (C-D) Dotplot analysis of select WNT signaling proteins in (C) 3D and 2D samples or (D) by cell cluster.

**Figure 5:**
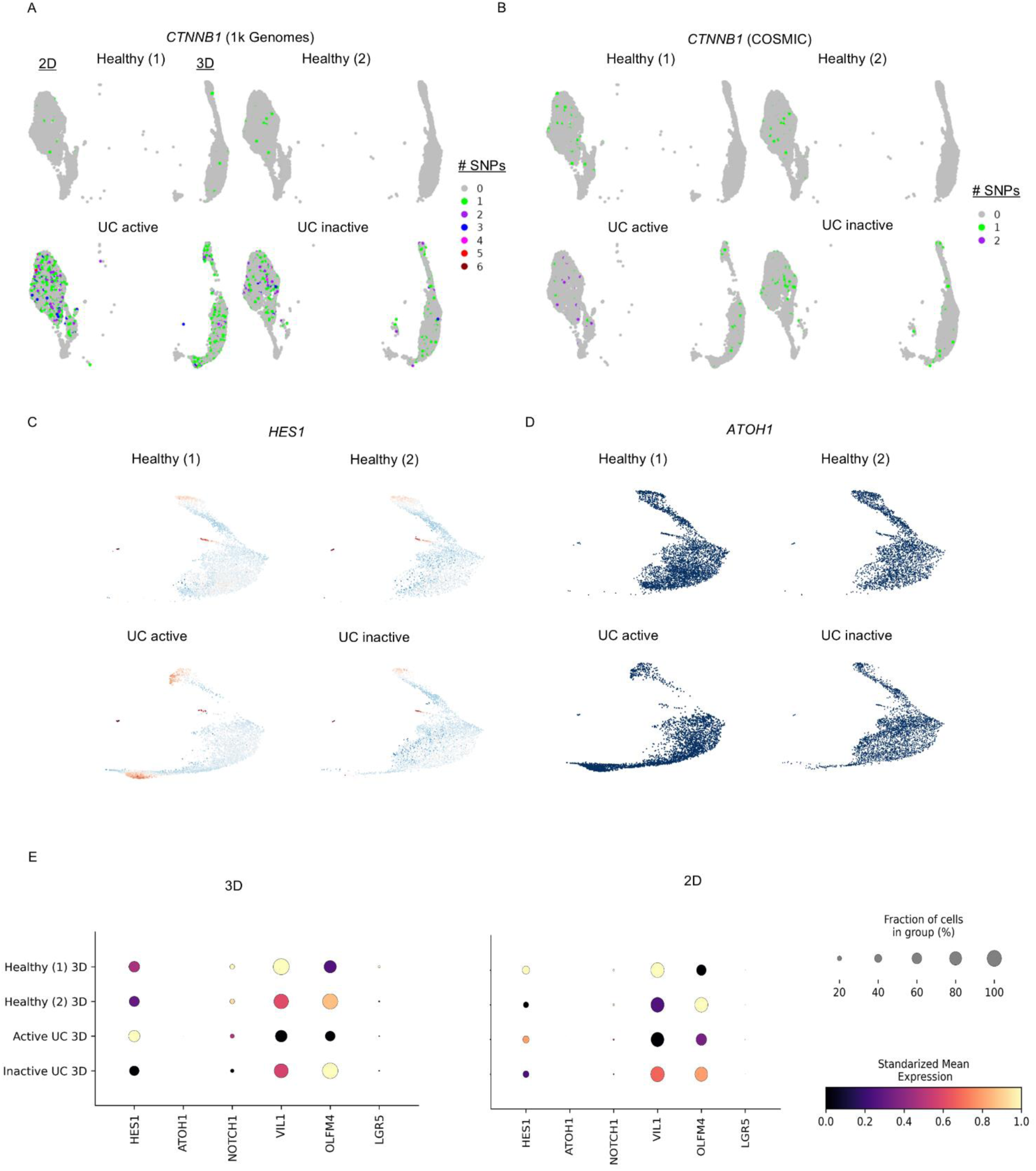
Aberrant WNT signaling in Active UC and downstream alterations. (A-B) SNPs found in single cell RNA sequencing using (A) 1k genomes database and (B) COSMIC database in 2D and 3D samples. UMAP analysis of gene expression for NOTCH signaling proteins (C) HES1 and (D) ATOH1 in 3D samples. (E) Dotplot analysis of select NOTCH signaling proteins as well as crypt top marker VIL1 and stem cell markers OLFM4 and LGR5 in 3D and 2D samples.

### NOTCH signaling is perturbed in UC colonoids

Given the altered WNT signaling observed in colonoids derived from active UC, we next investigated downstream effects on the NOTCH signaling pathway, which plays a critical role in regulating crypt-to-lumen epithelial differentiation (32). We focused on two transcriptional effectors of NOTCH signaling, *HES1* and *ATOH1*, which play opposing roles in regulating intestinal epithelial cell fate (32, 33). Our data revealed robust upregulation of HES1 in active UC colonoids, while ATOH1 expression was absent, consistent with prior observations in human enteroid models (34) (**Figure 5C-D**). These findings indicate that elevated HES1 expression in the 3D culture condition of active UC colonoids may be driven by increased WNT signaling, highlighting a potential interplay between WNT and NOTCH pathways (**Figure 5E**). As has been previously reported, *HES1* upregulation influences ISC differentiation, as well as stem cell replenishment (33, 35). In line with these findings, the active UC sample showed decreased expression of crypt-top marker *VIL1* and crypt ISC markers *LGR5* and *OLFM4* in both 3D and 2D culture conditions(**Figure 5E, S1**). Together, these findings suggest that upregulation of β-catenin and HES1 in active UC may contribute to impaired epithelial differentiation and maturation.

### Epithelial self-renewal is dysregulated in colonoids from individuals with active UC

RNAscope and immunostaining were employed on 2D and 3D colonoids to validate snRNA-seq and snATAC-seq findings by examining the spatial distribution of stem cell, crypt-top, and WNT signaling markers, including β-catenin. Additionally, these experiments were extended by including samples from an additional individual with active colonic inflammation (Active UC (1)) (**Table 1B**). As seen in **Figure 6A**, *OLFM4* expression is significantly reduced in both 3D (p < 0.0001) and 2D (p< 0.001) colonoids cultures derived from the two active UC samples, compared to those from healthy and inactive UC donors. This reduction in stem cell marker expression suggests selective depletion or dysfunction of the intestinal stem cell niche in colonoids derived from actively inflamed colonic mucosa. In addition, *VIL1* expression was significantly decreased (p < 0.001) in active UC colonoids compared to healthy and inactive UC colonoids, indicating impaired differentiation of mature epithelial cells (**Figure 6B**). Taken together, these findings indicate dysregulated self-renewal of colonic epithelial cells in active UC. To validate the RNA sequencing results, colonoids were immunostained for β-catenin expression. In 3D colonoids derived from active UC donors, both total β-catenin and active β-catenin (ABC) were statistically increased (p < 0.001) compared to those from healthy and inactive UC samples (**Figure 6C**). Interestingly, in 2D colonoids, total β-catenin expression remained comparable across groups. In healthy and inactive UC samples, β-catenin was predominantly localized to cell-cell junctions (**Figure 6B**). These findings support the conclusion that colonoids derived from actively inflamed UC mucosa exhibit enhanced WNT signaling through β-catenin, that is accompanied by impaired crypt luminal epithelial differentiation, likely due to intrinsic changes in stem cell maintenance and epithelial maturation.

**Figure 6:**
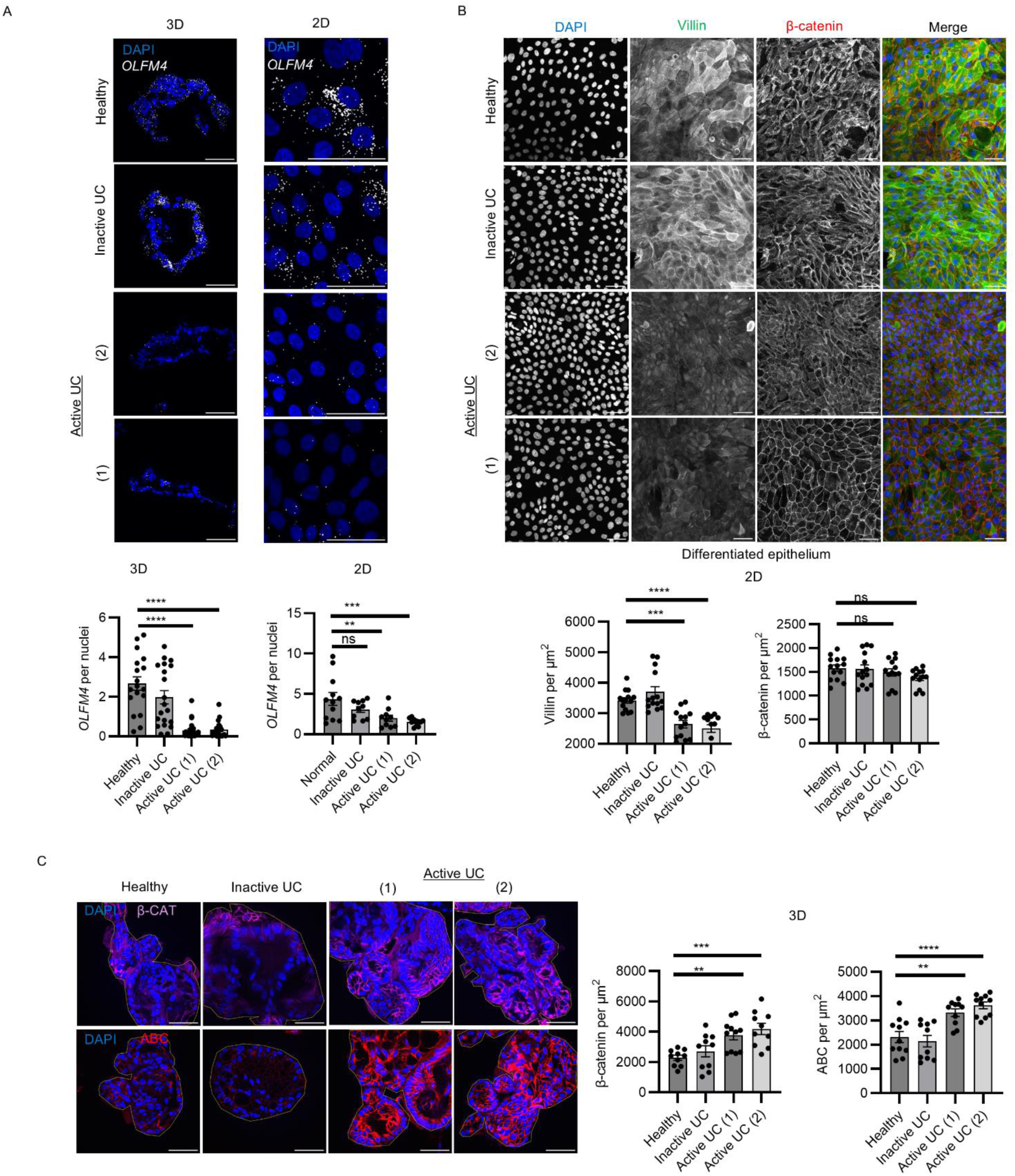
Active UC colonoids have dysregulated β-catenin and altered self-renewal. (A) In situ hybridization (RNAscope) of LGR5 and OLFM4 mRNA in 3D and 2D colonoids from various UC individuals (N=3 independent experiments). (B) Immunofluorescence of Villin and β-catenin in 2D colonoids from various UC individuals (N=3 independent experiments). (C) Immunofluorescence of total β-catenin (BCAT) and active β-catenin (ABC) in 3D colonoids from various UC individuals (N=3 independent experiments.

### Inhibition of RUNX2 restores self-renewal capacity in active UC colonoids

Given that RUNX2 mRNA and motifs accessibility were selectively enriched on active UC colonoids, we hypothesized that RUNX2 activity contributes to impaired epithelial self-renewal. To determine this, we treated 3D colonoid cultures with the small molecule RUNX2 inhibitor CADD522 (CADD) (36, 37). As shown in **Figure 7A**, treatment of active UC colonoids with CADD for 5 days resulted in a significant reduction on immunofluorescence labeling of both total (p<0.001) and active β-catenin (p<0.0001), indicating decreased WNT signaling. Furthermore, pre-treatment with CADD for 7 days prior to colonoid differentiation on collagen significantly increased Villin expression (p<0.01) and enhanced junctional localization of active β-catenin (p<0.05) in active UC colonoids (**Figure 7B**). Together, these findings suggest that aberrant WNT signaling in UC has an intrinsic epithelial component that impairs crypt luminal differentiation and is, at least in part, regulated by RUNX2 activity.

**Figure 7:**
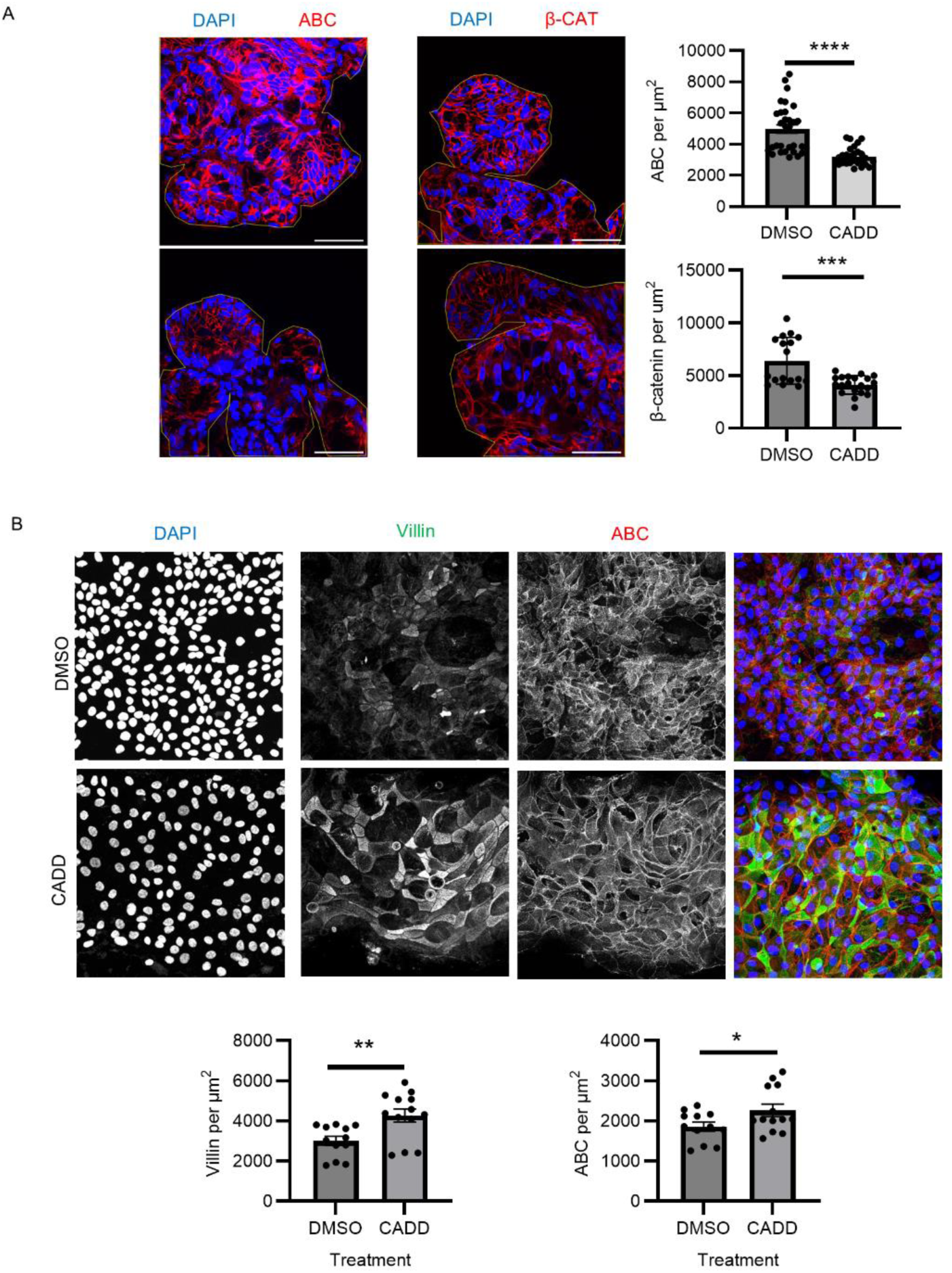
RUNX2 inhibition ameliorates the active UC self-renewal defect. (A) Immunofluorescence of β-catenin and active β-catenin (ABC) in 3D colonoids from various UC individuals treated with vehicle (DMSO) or 50 uM RUNX2 inhibitor (CADD) for 5 days (N=3 independent experiments). (B) Immunofluorescence of Villin and ABC in 2D colonoids from individual with severe active UC after 7-day pretreatment with CADD prior to 3-day differentiation (N=3 independent experiments).

## Discussion

Using patient derived human colonoids that faithfully recreate the crypt base-to-luminal axis, we uncovered inflammation-dependent, RUNX2-driven activation of WNT/β-catenin signaling that modulates epithelial self-renewal and differentiation in active UC. Specifically, a unique 3-D crypt-like cluster from active UC showed (i) elevated CTNNB1 and reduced APC, (ii) heightened NOTCH effector HES1 expression, and (iii) concomitant loss of the stem-cell marker OLFM4 and the enterocyte marker VIL1. Together these changes implicate RUNX2 as an epigenetic controller that amplifies WNT signaling and skews lineage decisions during intestinal inflammation.

### RUNX family epigenetics in UC, and colitis-associated colorectal cancer

RUNX transcription factors are lineage determining regulators that modulate gene expression by binding enhancer elements and recruiting chromatin modifiers such as p300/CBP, and HDAC complexes. In hematopoiesis, RUNX1 and RUNX3 coordinate sequential T- and NK-cell checkpoints (38–41), while in the gastrointestinal tract they enforce epithelial differentiation (42, 43). Importantly, the RUNX locus itself is a hotspot for epigenetic remodeling. Promoter hyper-methylation silences RUNX3 in gastric and colorectal cancer, whereas enhancer hijacking and super-enhancer formation have been described for RUNX2 in osteosarcoma (25, 44). Such plasticity makes the RUNX family sensitive to inflammatory cytokines and metabolic stress as observed in UC.

Our analysis of single-cell databases (4, 26) reveals an induction of all three RUNX proteins in inflamed epithelial cells. This observation aligns with in vivo evidence showing that conditional Runx1 deletion within intestinal epithelial cells (IECs) triggers spontaneous tumors in mice (45), and that deletion of Runx3 in leukocytes results in colonic crypt hyperplasia and i inflammation (46). Taken together data highlight the protective role of RUNX on both epithelial and immune cells. Mechanistically, RUNX transcription factors can both potentiate and restrain WNT signaling. For example, RUNX2 enhances transcription of EMT genes in colon cancer cells with high WNT activity (47, 48). Indeed, in Crohn’s disease tissue RUNX2 up regulates p53 and limits apoptosis (49), whereas in UC specific clusters co-expression of RUNX2 with ANXA5, a pro-apoptotic effector, was observed highlighting context-specific toggling between protective and pathological outcomes (50).

The implications for colitis-associated colorectal cancer (CA-CRC) are significant. Classic sporadic CRC follows the APC-KRAS-p53 mutation sequence, but in CA-CRC, APC mutations are comparatively rare (51). Therefore, it will be interesting to further investigate if sustained RUNX2/β-catenin signaling could substitute for APC loss, gradually tipping epithelial progenitors toward dysplasia. These nuances suggest that selective modulation of RUNX activity may be required to restore epithelial equilibrium without unleashing oncogenic programs. Therefore exploring small-molecule inhibitors that discriminate among RUNX paralogs (e.g., CADD522), which is currently being developed for potential cancer therapeutics, to treat inflamed epithelial cells and prevent the development of CA-CRC represents a therapeutic avenue worthy of pre-clinical evaluation (36, 37).

#### Crosstalk between WNT and NOTCH in epithelial differentiation

WNT and NOTCH pathways form a tightly interlaced network that orchestrates the balance between stem-cell maintenance, lineage commitment, and barrier integrity along the crypt-luminal axis (52). Canonical WNT signaling, driven by β-catenin accumulation, is indispensable for crypt base stem cell proliferation, whereas NOTCH signaling modulates the binary decision between secretory and absorptive progenitors, largely through HES family transcriptional repressors (33, 52). Our human colonoid model, cultured in high-WNT medium and devoid of immune/mesenchymal partners, isolates the epithelial-intrinsic layer of this process.

Within this controlled environment we observe an intriguing dissociation. β-catenin targets (CTNNB1, AXIN2) along with Hes1 are up-regulated, yet the archetypal stem marker Olfm4 is suppressed in active UC states. This contrasts with tissue biopsies where OLFM4 marks crypt base stem cells and often expands in UC (26, 53). Two non-exclusive mechanisms could explain the discrepancy. First, immune cell derived cytokines such as IL-22 and IL-8, abundant in vivo and known to boost OLFM4 transcription, are absent in organoids (53). Second, our data hint at a shift from rapidly cycling LGR5⁺ crypt base stem cells to +4 slow-cycling stem cells (BMI1⁺ LGR5⁻), a population favored when NOTCH signaling is perturbed (18).

The pervasive expression of MUC2 in our cultures could also be important to consider. Goblet cell expansion is often viewed as compensatory in UC, enhancing the mucus barrier; yet studies indicate that while goblet cells are vital for luminal defense, they are dispensable for crypt regeneration (32). Our data suggest that during active inflammation, epithelial cells attempt to maintain barrier function through MUC2 while reallocating stem-cell output toward absorptive lineages. RUNX2 may act as a molecular hub in this reprogramming. β-catenin can bind to the RUNX2 promoter, and RUNX2, in turn, can trans-activate HES1, linking WNT levels to NOTCH expression.

## Conclusion

Our data position RUNX2 as a pivotal epigenetic switch that amplifies WNT/β-catenin and propagates NOTCH activity during intestinal inflammation, but the precise molecular mechanism remains undefined. Therapeutically, small molecule interference with RUNX2 offers an attractive strategy to restore epithelial barrier function in UC. The RUNX2 antagonist CADD522, currently being evaluated for cancer therapy (36, 37) could be repurposed to modulate WNT signaling and restore adequate stem cell differentiation in Ulcerative Colitis.

In summary, we demonstrate an inflammation-dependent RUNX2-dependent mechanism that activates WNT/β-catenin signaling and contributes to dysregulation of intestinal epithelial self-renewal. Increased understanding of regulation of WNT/ β-catenin signaling can identify novel targets to inform rational therapies for Ulcerative Colitis in the future.

## Methods

### Sex as a biological variable

Our study examined de-identified human tissues from both male and female donors, and similar findings are reported for both sexes.

### Reagents

Anti-β-catenin (IF:1/300), anti-ABC (EMD Millipore, Clone 8E7, Cat. 05-665; IF: 1/300), anti-Villin (Genetex, Cat. GTX109940; IF: 1/200), anti-MUC2 (Abcam, Cat. ab90007; IF:1/200) antibodies were purchased from corresponding vendors. Alexa Fluor 488-conjugated donkey anti-rabbit (IF:1/1000) and Alexa Fluor 555-conjugated donkey-anti-mouse (IF:1/1000) secondary antibody were purchased from Thermofisher Scientific. RUNX2 inhibitor CADD522 (Cat. 36679) were purchased from Cayman Chemical and used at a concentration of 100 µM.

### Human subjects

Normal, de-identified human colonoids were purchased from the translational tissue modeling laboratory (TTML) at the University of Michigan. All human tissue used in this work was de-identified.

### Human colonoids

Human colonoids were provided by the University of Michigan Translational Tissue Modeling Laboratory (TTML) and maintained in Matrigel (BD Biosciences) in GMGF media composed of Advanced DMEM/F12, 10% FBS, 2 mm GlutaMAX-1, 10 mm HEPES, 100 U/ml Penicillin/streptomycin (all from Invitrogen) and L-WRN conditioned media (54).

2D colonoid monolayers were prepared as previously described (55). In brief, human 3D colonoid single-cell suspension was obtained by resuspension in 0.05% Trypsin/0.5 mm EDTA and vigorously pipetting up and down. Trypsin was inactivated by adding 1 ml advanced DMEM/F12- containing 10% FBS. Dissociated cells were passed through a 40 µm cell strainer, then cultured in collagen type IV-coated 48-well tissue culture plates (72 h). After an initial 24 hours in LWRN-containing media, 2D cultures were maintained in differentiation media containing GMGF media supplemented with 10 nm Gastrin and 1 mm *N-*Acetylcysteine (Sigma), 10 mm Nicotinamide, 10 µm SB202190 (StemCell Tech), 500 nm A83-01 (StemCell Tech), 10 ng/ml EGF (R&D systems).

### snRNA/snATAC multiomic sequencing

The colonoid experiments were carefully conducted in two separate batches to reduce batch effects in scRNA-seq and snATAC-seq data. One batch with healthy (1) and active UC were carried out in parallel for 2D and 3D conditions and experiments for healthy (2) and inactive UC were carried out in a subsequent batch. Cells were harvested and dissociated into single cell suspensions in parallel using TRYPLE (Gibco, cat 50-591-419), frozen at -80°C, then thawed and prepared for single nuclei suspension for the 10X Chromium system (see below). Since the 10X Chromium system allows parallel processing of multiple samples at a time, cells were captured (Gel bead-in-Emulsion - GEMS) and processed (i.e., library prep) in parallel. Batch samples were sequenced together on a Novaseq 6000.

For analyzing snRNA sequencing from these individuals, we processed samples by thresholding for optimal RNA yield per cell (RNA between 1000-50000, ATAC between 1000-100000, UMI>200, mitochondrial genes <30%), normalized and scaled using SCTransform (with regression including percentage of mitochondria genes, UMI, and cell cycle genes), performed PCA for linear dimensional reduction, clustered cells based on K-nearest neighbor, and finally performed UMAP non-linear dimensional reduction to visual similar cells in a low dimensional space. Samples were then merged based on 3D and 2D samples and integrated together using Seurat’s CCA integration method and reclustered using Seurat FindClusters (56).

For analyzing snATAC sequencing, we processed samples using ArchR (57) wherein cells were thresholded for TSS>5, doublets were excluded, 500 bp tiles were obtained through iterative LSI, clustered using Seurat FindClusters, and embedded onto a UMAP. Samples were then integrated using Harmony separately for 3D and 2D samples and reclustered using Seurat FindClusters. Pseudobulk replicates were obtained for a maximum of 4 samples per cluster and Peak Calling was performed on each cluster using MACS2 (58). Motifs were identified based on the corresponding peaks using CIS-BP motif set.

Pseudotime analysis was performed using the Python package CellRank2 (59). Briefly, raw counts were imputed using Markov Affinity-based Graph Imputation of Cells (MAGIC) before using the Palantir algorithm to identify pseudotime trajectories. Afterwards, we utilize the “PseudotimeKernel” from CellRank2 to compute the transition matrix and plot the trajectories projections. To obtain unbiased initial and terminal states in the pseudotime transition matrix, we performed Generalized Perron Cluster Cluster Analysis (GPCCA) to identify macrostates of cellular dynamics. Then, we visualize a coarse-grained transition matrix among the macrostates to classify them into initial and terminal states.

For PseudoBulk RNA-seq we utilize the Seurat AggregateExpression function. DEG analysis was performed using the DESeq2 R package (60). GSEA analysis was performed using the GSEA function on the clusterProfiler R package. Volcano plots were done using the EnhancedVolcano R package. DotPlots and UMAP plots were performed using the plotting function of Seurat or Scanpy in R or python, respectively. GSEA plots were performed with the ggplot2 package, and gene trends plots were performed with the Palantir python package.

To identify SNPs in the scRNA-seq FASTQ files, we used the Cellsnp-lite program (61). Briefly, we used the post-sorted BAM files generated by the Cell Ranger pipeline during alignment. BAM files were compared to the 1K Genome (29) and COSMIC SNPs databases (30). Barcodes were provided to track SNPs in each cell. Cellsnp-lite was used in its “1a Mode” protocol with a minimum read count of 20 and a minimum minor allele frequency of 0.01. Cellsnp-lite output results were processed in R and added to scRNA-seq metadata for visualization on UMAP plots.

### Immunofluorescence and RNAscope

2D colonoids monolayers were grown on plastic chamber slides (Thermo Fisher Scientific, Cat. 177445) and fixed with 4% paraformaldehyde (PFA). PFA-fixed monolayers were permeabilized with 0.5% Triton X-100 for 10 minutes. Monolayers were blocked with 3% donkey serum in DPBS with 0.05% Tween-20 blocking buffer for 1 hour. Primary antibodies were diluted in blocking buffer, and cells were incubated overnight at 4°C. Cells were washed with PBS with 0.05% Tween 20, and fluorescently labeled secondary antibodies were diluted in blocking buffer followed by incubation for 1 hour at room temperature. Cells were washed and mounted in Prolong Gold antifade agent (Thermo Fisher Scientific, P36930).

3D colonoids were recovered in Cell Recovery Solution (Corning) and fixed in 4% PFA for 1 hour. Colonoids were then permeabilized with 1% Triton X-100 for 1 hour, blocked with 3% BSA in DPBS with 0.05% Tween-20 blocking buffer for 1 hour. Primary antibodies were diluted in blocking buffer and cells were incubated overnight at 4°C. After washing, cells were then blocked in secondary antibodies diluted in blocking buffer overnight at 4°C. Cells were then washed and mounted in Prolong Gold antifade agent. Localization of human *LGR5/OLFM4* mRNA was determined using whole 2D monolayers or cryosections of 3D colonoids after embedding in OCT with formalin-fixation and ethanol-dehydration. Following incubation with hydrogen peroxide (ACDbio, Cat. 322000), tissues were digested by protease III treatment (Cat. 322330). *LGR5/OLFM4* probe for human (ACDbio, Cat. No. 311021, Cat. Cat 311041-C2) was hybridized for 2 h followed by signal amplification.

Fluorescence imaging was performed with a Nikon A1 confocal microscope (Nikon) in the Microscopy & Image Analysis Laboratory Core at the University of Michigan and analyzed using ImageJ.

### Statistics

Parametric or nonparametric tests were performed to evaluate statistical significance after finding if datasets were normally distributed for immunofluorescence. Statistical significance was measured by Student’s t test, one-way or two-way ANOVA with indicated multiple comparisons test using Graphpad Prism software. Significance was set as p ≤ 0.05. Results are expressed as means ± standard error of mean.

## Data availability

The authors declare that all data supporting the findings of this study are available within the paper and its supplementary information files, or from the corresponding author on reasonable request. Multi-omic sequencing data generated in this study has been deposited at EMBL-EBI ArrayExpress. The accession number will be provided upon publication

## Code availability

All code used in this project will be deposited in a public GitHub repository; the link will be released following publication.

## Author Contributions

ZSW & RICS planned and were involved in all experiments as well as wrote the manuscript. Order of appearance was agreed by ZSW & RICS. JM prepared the cells for all experiments involving immunofluorescence and in situ hybridization (RNAscope). SF was involved in the immunofluorescence of the 2D human colonoids. JCB, AN, and CP were involved in the preparation of the manuscript. SB provided the active UC human colonoid line HT404 (Table 1B). JC was involved in the data analysis for ATAC- and RNA-sequencing as well as the preparation of the manuscript.

## Acknowledgements

We thank the Advanced Genomics Core and the TTML at the University of Michigan Medical School. The authors additionally thank the laboratory of Mary Estes at Baylor University for assistance in enteroid culture generation as well as Chithra Kannachazhath Muraleedharan for technical assistance. This work was supported by the following NIH grants: R01DK055679 and R01DK059888 (AN), T32HL007517 (ZSW), R56DK140172 (JCB), R01DK129058, R01DK129214, and R01DK079392 (CAP).

## References

1. Blander JM. Death in the intestinal epithelium-basic biology and implications for inflammatory bowel disease. FEBS J. 2016;283(14):2720–30.

2. Javaid SS, Akhtar S, Hafeez A, Nofal A, Rahman S, Farooqui SK, et al. Trends in Mortality Due to Inflammatory Bowel Disease in the United States: A CDC WONDER Database Analysis (1999-2020). Dig Dis Sci. 2025;70(2):494–503.

3. Kaplan GG. The global burden of IBD: from 2015 to 2025. Nat Rev Gastroenterol Hepatol. 2015;12(12):720–7.

4. Murthy S, Anbazhagan M, Maddipatla SC, Kolachala VL, Dodd A, Pelia R, et al. Single-cell transcriptomics of rectal organoids from individuals with perianal fistulizing Crohn’s disease reveals patient-specific signatures. Sci Rep. 2024;14(1):26142.

5. van der Flier LG, and Clevers H. Stem cells, self-renewal, and differentiation in the intestinal epithelium. Annu Rev Physiol. 2009;71:241–60.

6. Cosin-Roger J, Ortiz-Masia D, Calatayud S, Hernandez C, Alvarez A, Hinojosa J, et al. M2 macrophages activate WNT signaling pathway in epithelial cells: relevance in ulcerative colitis. PLoS One. 2013;8(10):e78128.

7. Cosin-Roger J, Ortiz-Masia D, Calatayud S, Hernandez C, Esplugues JV, and Barrachina MD. The activation of Wnt signaling by a STAT6-dependent macrophage phenotype promotes mucosal repair in murine IBD. Mucosal Immunol. 2016;9(4):986–98.

8. Cosin-Roger J, Ortiz-Masia MD, and Barrachina MD. Macrophages as an Emerging Source of Wnt Ligands: Relevance in Mucosal Integrity. Front Immunol. 2019;10:2297.

9. Macias-Ceja DC, Coll S, Bauset C, Seco-Cervera M, Gisbert-Ferrandiz L, Navarro F, et al. IFNgamma-Treated Macrophages Induce EMT through the WNT Pathway: Relevance in Crohn’s Disease. Biomedicines. 2022;10(5).

10. Raveenthiraraj S, Awanis G, Chieppa M, O’Connell AE, and Sobolewski A. M1 and M2 Macrophages Differentially Regulate Colonic Crypt Renewal. Inflamm Bowel Dis. 2024;30(7):1138–50.

11. Deng L, He XC, Chen S, Zhang N, Deng F, Scott A, et al. Frizzled5 controls murine intestinal epithelial cell plasticity through organization of chromatin accessibility. Dev Cell. 2025;60(3):352–63 e6.

12. Ray G, and Longworth MS. Epigenetics, DNA Organization, and Inflammatory Bowel Disease. Inflamm Bowel Dis. 2019;25(2):235–47.

13. Xu J, Xu HM, Yang MF, Liang YJ, Peng QZ, Zhang Y, et al. New Insights Into the Epigenetic Regulation of Inflammatory Bowel Disease. Front Pharmacol. 2022;13:813659.

14. Zou WY, Blutt SE, Crawford SE, Ettayebi K, Zeng XL, Saxena K, et al. Human Intestinal Enteroids: New Models to Study Gastrointestinal Virus Infections. Methods Mol Biol. 2019;1576:229–47.

15. Huang L, Xu Z, Lei X, Huang Y, Tu S, Xu L, et al. Paneth cell-derived iNOS is required to maintain homeostasis in the intestinal stem cell niche. J Transl Med. 2023;21(1):852.

16. Jaitner S, Pretzsch E, Neumann J, Schaffauer A, Schiemann M, Angele M, et al. Olfactomedin 4 associates with expression of differentiation markers but not with properties of cancer stemness, EMT nor metastatic spread in colorectal cancer. J Pathol Clin Res. 2023;9(1):73–85.

17. Li H, Chaitankar V, Zhu J, Chin K, Liu W, Pirooznia M, et al. Olfactomedin 4 mediation of prostate stem/progenitor-like cell proliferation and differentiation via MYC. Sci Rep. 2020;10(1):21924.

18. Srinivasan T, Than EB, Bu P, Tung KL, Chen KY, Augenlicht L, et al. Notch signalling regulates asymmetric division and inter-conversion between lgr5 and bmi1 expressing intestinal stem cells. Sci Rep. 2016;6:26069.

19. Yan KS, Chia LA, Li X, Ootani A, Su J, Lee JY, et al. The intestinal stem cell markers Bmi1 and Lgr5 identify two functionally distinct populations. Proc Natl Acad Sci U S A. 2012;109(2):466–71.

20. Zhu Y, Huang YF, Kek C, and Bulavin DV. Apoptosis differently affects lineage tracing of Lgr5 and Bmi1 intestinal stem cell populations. Cell Stem Cell. 2013;12(3):298–303.

21. de Lau W, Kujala P, Schneeberger K, Middendorp S, Li VS, Barker N, et al. Peyer’s patch M cells derived from Lgr5(+) stem cells require SpiB and are induced by RankL in cultured “miniguts". Mol Cell Biol. 2012;32(18):3639–47.

22. Wang B, Iglesias-Ledon L, Bishop M, Chadha A, Rudolph SE, Longo BN, et al. Impact of Micro-and Nano-Plastics on Human Intestinal Organoid-Derived Epithelium. Curr Protoc. 2024;4(4):e1027.

23. Korsunsky I, Millard N, Fan J, Slowikowski K, Zhang F, Wei K, et al. Fast, sensitive and accurate integration of single-cell data with Harmony. Nat Methods. 2019;16(12):1289–96.

24. Lasconi C, Pahl MC, Cousminer DL, Doege CA, Chesi A, Hodge KM, et al. Variant-to-Gene-Mapping Analyses Reveal a Role for the Hypothalamus in Genetic Susceptibility to Inflammatory Bowel Disease. Cell Mol Gastroenterol Hepatol. 2021;11(3):667–82.

25. Vega OA, Lucero CMJ, Araya HF, Jerez S, Tapia JC, Antonelli M, et al. Wnt/beta-Catenin Signaling Activates Expression of the Bone-Related Transcription Factor RUNX2 in Select Human Osteosarcoma Cell Types. J Cell Biochem. 2017;118(11):3662–74.

26. Smillie CS, Biton M, Ordovas-Montanes J, Sullivan KM, Burgin G, Graham DB, et al. Intra- and Inter-cellular Rewiring of the Human Colon during Ulcerative Colitis. Cell. 2019;178(3):714–30 e22.

27. Liu J, Xiao Q, Xiao J, Niu C, Li Y, Zhang X, et al. Wnt/beta-catenin signalling: function, biological mechanisms, and therapeutic opportunities. Signal Transduct Target Ther. 2022;7(1):3.

28. Koch S. Extrinsic control of Wnt signaling in the intestine. Differentiation. 2017;97:1–8.

29. Genomes Project C, Auton A, Brooks LD, Durbin RM, Garrison EP, Kang HM, et al. A global reference for human genetic variation. Nature. 2015;526(7571):68–74.

30. Sondka Z, Dhir NB, Carvalho-Silva D, Jupe S, Madhumita, McLaren K, et al. COSMIC: a curated database of somatic variants and clinical data for cancer. Nucleic Acids Res. 2024;52(D1):D1210–D7.

31. Khera AV, Chaffin M, Aragam KG, Haas ME, Roselli C, Choi SH, et al. Genome-wide polygenic scores for common diseases identify individuals with risk equivalent to monogenic mutations. Nat Genet. 2018;50(9):1219–24.

32. Banjac I, Maimets M, Tsang IHC, Dioli M, Hansen SL, Krizic K, et al. Fate mapping in mouse demonstrates early secretory differentiation directly from Lgr5+ intestinal stem cells. Dev Cell. 2025.

33. VanDussen KL, Carulli AJ, Keeley TM, Patel SR, Puthoff BJ, Magness ST, et al. Notch signaling modulates proliferation and differentiation of intestinal crypt base columnar stem cells. Development. 2012;139(3):488–97.

34. Farhadipour M, Arnauts K, Clarysse M, Thijs T, Liszt K, Van der Schueren B, et al. SCFAs switch stem cell fate through HDAC inhibition to improve barrier integrity in 3D intestinal organoids from patients with obesity. iScience. 2023;26(12):108517.

35. Fre S, Huyghe M, Mourikis P, Robine S, Louvard D, and Artavanis-Tsakonas S. Notch signals control the fate of immature progenitor cells in the intestine. Nature. 2005;435(7044):964–8.

36. Green D, Singh A, Tippett VL, Tattersall L, Shah KM, Siachisumo C, et al. YBX1-interacting small RNAs and RUNX2 can be blocked in primary bone cancer using CADD522. J Bone Oncol. 2023;39:100474.

37. Kim MS, Gernapudi R, Choi EY, Lapidus RG, and Passaniti A. Characterization of CADD522, a small molecule that inhibits RUNX2-DNA binding and exhibits antitumor activity. Oncotarget. 2017;8(41):70916–40.

38. de Bruijn M, and Dzierzak E. Runx transcription factors in the development and function of the definitive hematopoietic system. Blood. 2017;129(15):2061–9.

39. Ohno S, Sato T, Kohu K, Takeda K, Okumura K, Satake M, et al. Runx proteins are involved in regulation of CD122, Ly49 family and IFN-gamma expression during NK cell differentiation. Int Immunol. 2008;20(1):71–9.

40. Seo W, and Taniuchi I. The Roles of RUNX Family Proteins in Development of Immune Cells. Mol Cells. 2020;43(2):107–13.

41. Dybska E, Adams AT, Duclaux-Loras R, Walkowiak J, and Nowak JK. Waiting in the wings: RUNX3 reveals hidden depths of immune regulation with potential implications for inflammatory bowel disease. Scand J Immunol. 2021;93(5):e13025.

42. Peng W, Zeng C, Xu J, Zhao H, Zhu Q, Xu H, et al. Regulation of epithelial cell differentiation by the Ubiquitous expressed transcript isoform 1 in ulcerative colitis. J Gastroenterol Hepatol. 2023;38(11):2006–17.

43. Buchert M, Darido C, Lagerqvist E, Sedello A, Cazevieille C, Buchholz F, et al. The symplekin/ZONAB complex inhibits intestinal cell differentiation by the repression of AML1/Runx1. Gastroenterology. 2009;137(1):156–64, 64 e1-3.

44. Dutta B, and Osato M. The RUNX Family, a Novel Multifaceted Guardian of the Genome. Cells. 2023;12(2).

45. Fijneman RJ, Anderson RA, Richards E, Liu J, Tijssen M, Meijer GA, et al. Runx1 is a tumor suppressor gene in the mouse gastrointestinal tract. Cancer Sci. 2012;103(3):593–9.

46. Brenner O, Levanon D, Negreanu V, Golubkov O, Fainaru O, Woolf E, et al. Loss of Runx3 function in leukocytes is associated with spontaneously developed colitis and gastric mucosal hyperplasia. Proc Natl Acad Sci U S A. 2004;101(45):16016–21.

47. Verhagen MP, Xu T, Stabile R, Joosten R, Tucci FA, van Royen M, et al. The SW480 cell line as a model of resident and migrating colon cancer stem cells. iScience. 2024;27(9):110658.

48. Yi H, Li G, Long Y, Liang W, Cui H, Zhang B, et al. Integrative multi-omics analysis of a colon cancer cell line with heterogeneous Wnt activity revealed RUNX2 as an epigenetic regulator of EMT. Oncogene. 2020;39(28):5152–64.

49. Gu L, Zhao J, Zhang S, Xu W, Ni R, and Liu X. Runt-related transcription factor 2 (RUNX2) inhibits apoptosis of intestinal epithelial cells in Crohn’s disease. Pathol Res Pract. 2018;214(2):245–52.

50. Sweeney K, Cameron ER, and Blyth K. Complex Interplay between the RUNX Transcription Factors and Wnt/beta-Catenin Pathway in Cancer: A Tango in the Night. Mol Cells. 2020;43(2):188–97.

51. Rajamaki K, Taira A, Katainen R, Valimaki N, Kuosmanen A, Plaketti RM, et al. Genetic and Epigenetic Characteristics of Inflammatory Bowel Disease-Associated Colorectal Cancer. Gastroenterology. 2021;161(2):592–607.

52. Nakamura T, Tsuchiya K, and Watanabe M. Crosstalk between Wnt and Notch signaling in intestinal epithelial cell fate decision. J Gastroenterol. 2007;42(9):705–10.

53. Gersemann M, Becker S, Nuding S, Antoni L, Ott G, Fritz P, et al. Olfactomedin-4 is a glycoprotein secreted into mucus in active IBD. J Crohns Colitis. 2012;6(4):425–34.

54. Dame MK, Attili D, McClintock SD, Dedhia PH, Ouillette P, Hardt O, et al. Identification, isolation and characterization of human LGR5-positive colon adenoma cells. Development. 2018;145(6).

55. Reed M, Luissint AC, Azcutia V, Fan S, O’Leary MN, Quiros M, et al. Epithelial CD47 is critical for mucosal repair in the murine intestine in vivo. Nat Commun. 2019;10(1):5004.

56. Hao Y, Hao S, Andersen-Nissen E, Mauck WM, 3rd, Zheng S, Butler A, et al. Integrated analysis of multimodal single-cell data. Cell. 2021;184(13):3573–87 e29.

57. Granja JM, Corces MR, Pierce SE, Bagdatli ST, Choudhry H, Chang HY, et al. ArchR is a scalable software package for integrative single-cell chromatin accessibility analysis. Nat Genet. 2021;53(3):403–11.

58. Zhang Y, Liu T, Meyer CA, Eeckhoute J, Johnson DS, Bernstein BE, et al. Model-based analysis of ChIP-Seq (MACS). Genome Biol. 2008;9(9):R137.

59. Weiler P, Lange M, Klein M, Pe’er D, and Theis F. CellRank 2: unified fate mapping in multiview single-cell data. Nat Methods. 2024;21(7):1196–205.

60. Love MI, Huber W, and Anders S. Moderated estimation of fold change and dispersion for RNA-seq data with DESeq2. Genome Biol. 2014;15(12):550.

61. Huang X, and Huang Y. Cellsnp-lite: an efficient tool for genotyping single cells. Bioinformatics. 2021;37(23):4569–71.

