## Supplementary Figures for "RUNX2 promotes epigenetic WNT signaling in inflamed intestinal epithelial cells"

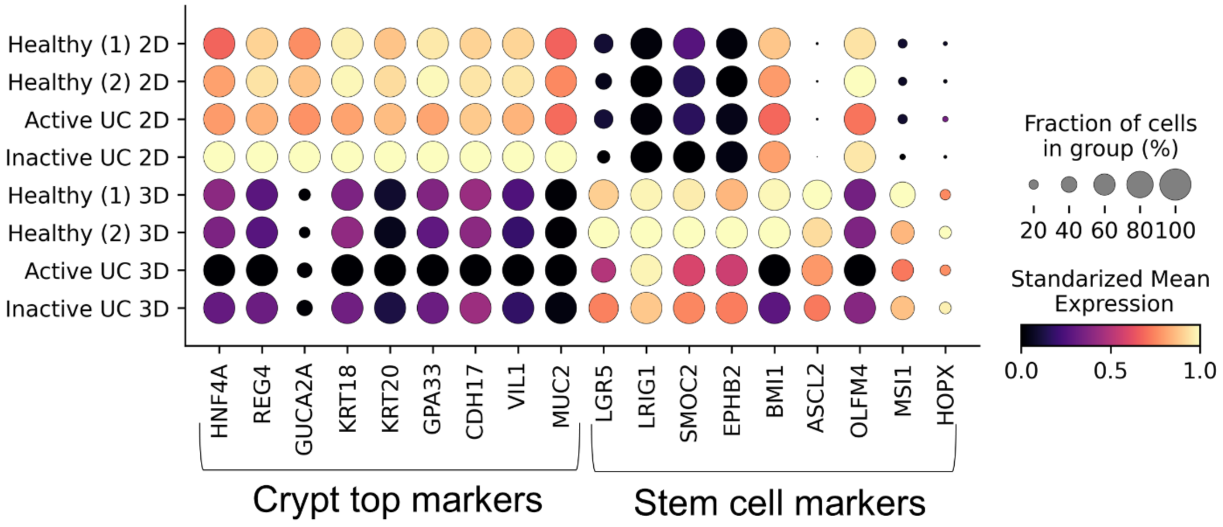


**Supplementary Figure 1.** Dot plot comparing the RNA expression of crypt-top (differentiated) and stem-cell (undifferentiated) marker genes across all 3D and 2D colonoid samples.

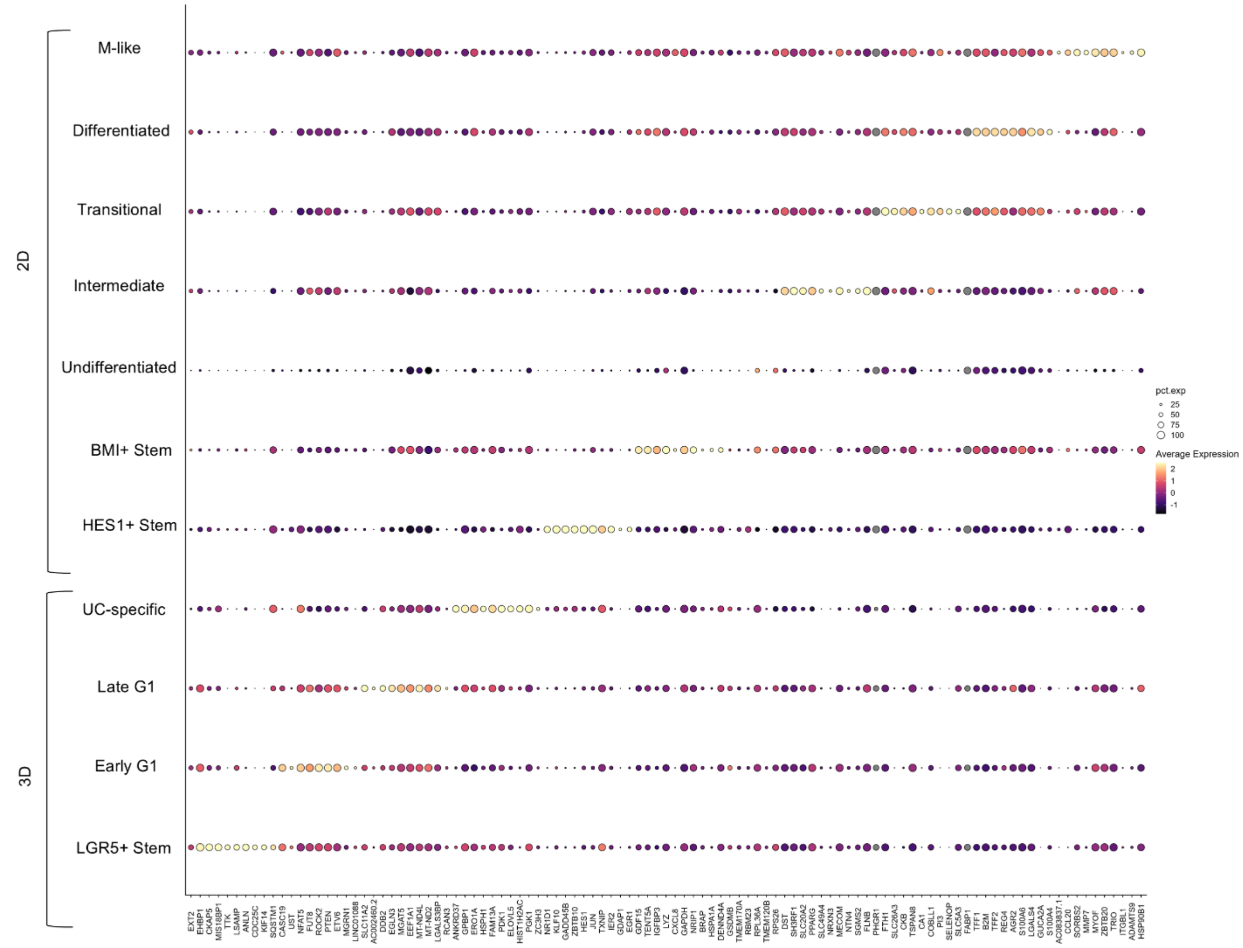


**Supplementary Figure 2.** Dot plot showing RNA expression of top 10 marker genes for each cluster on 3D and 2D colonoid samples.

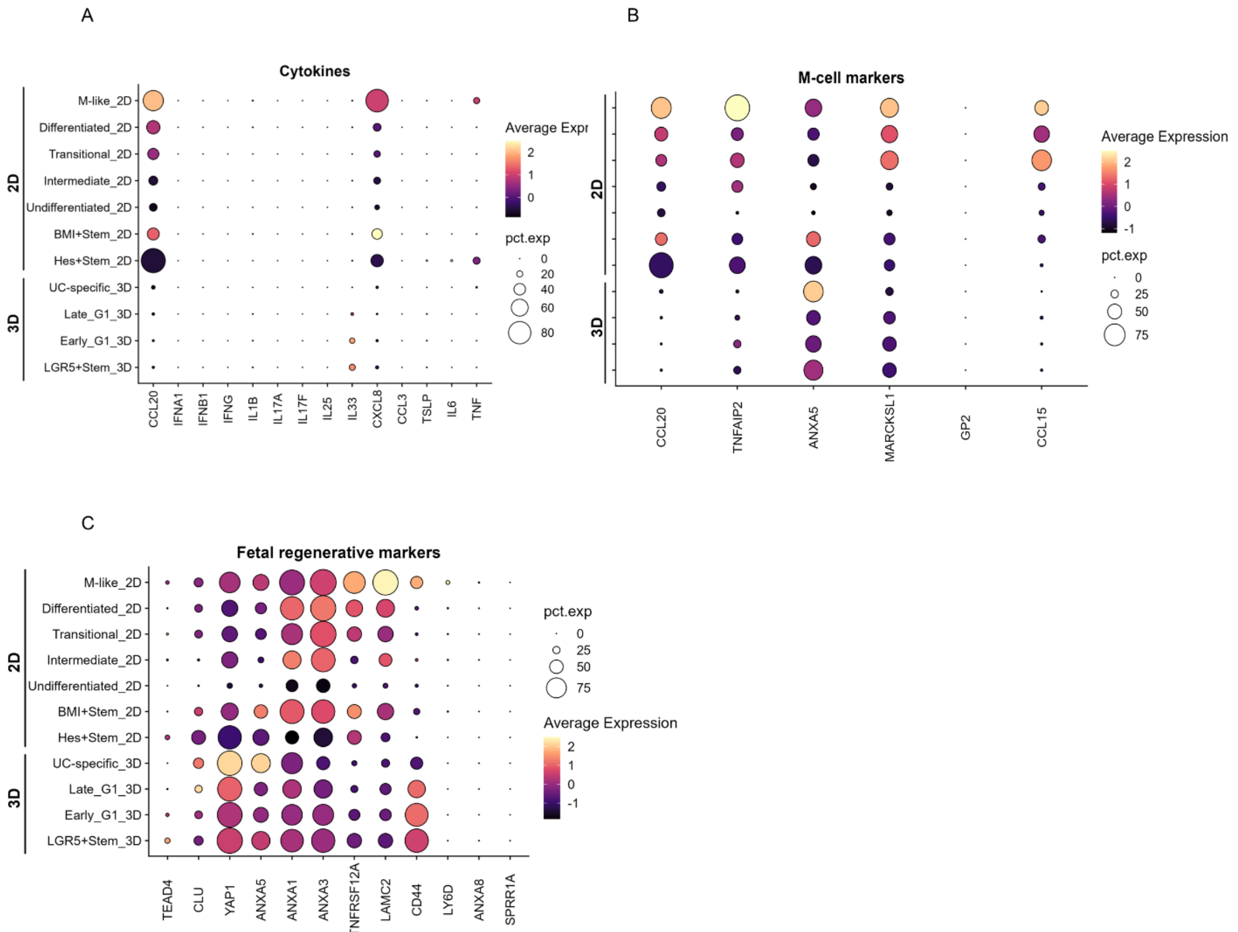


**Supplementary Figure 3.** Dot plots showing RNA expression associated with A) Cytokines, B) M-cell markers and C) Fetal regenerative markers across all clusters on 3D and 2D colonoid samples.

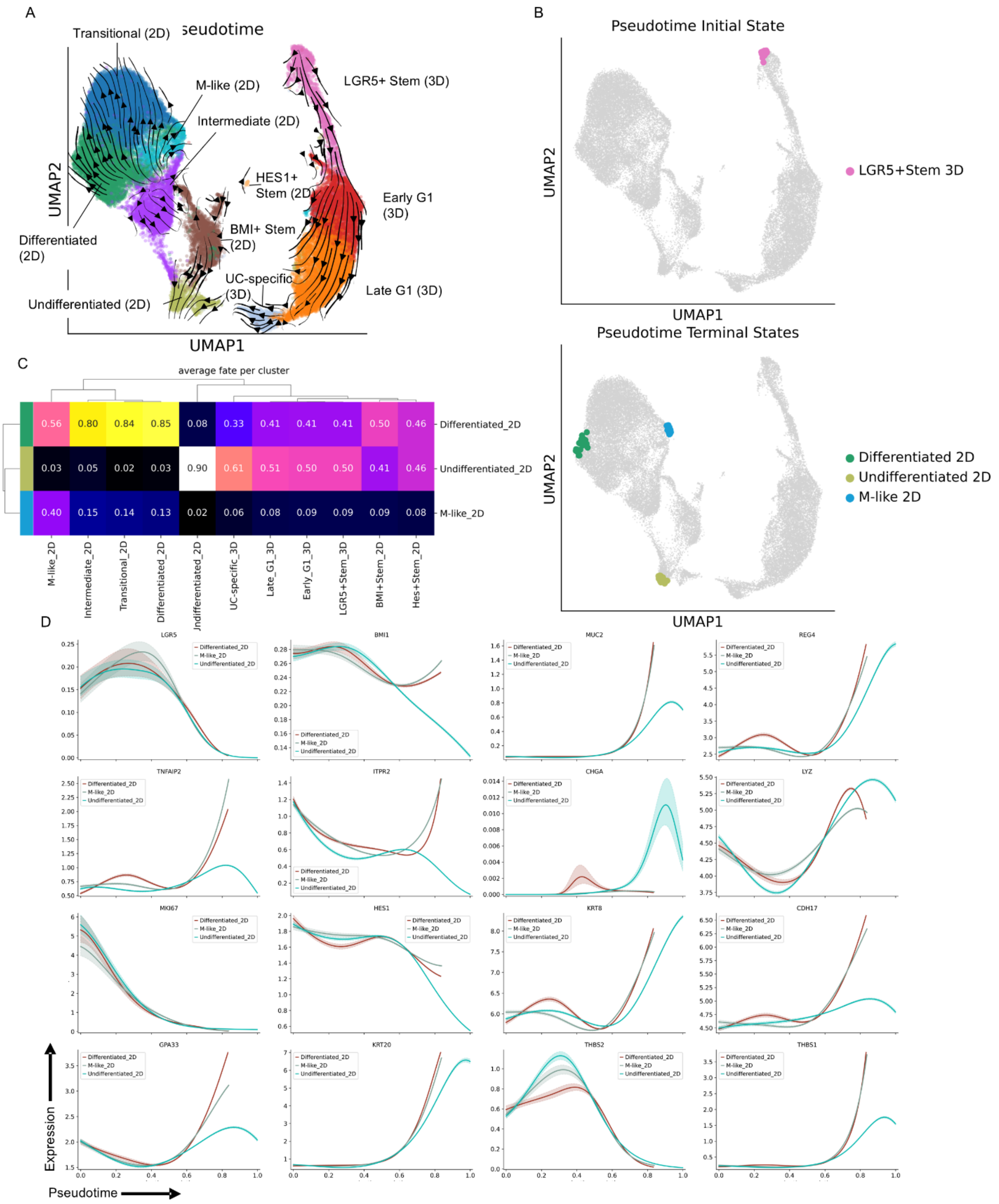


**Supplementary Figure 4.** A) UMAP of pseudotime transitions on RNA-seq of all 2D and 3D colonoid samples. B) UMAP with unbiased CellRank2-inferred initial and terminal states. C) Heatmap of fate probabilities for each cell type toward each terminal state. D) Pseudotime-aligned expression dynamics of marker genes along each terminal state trajectory.

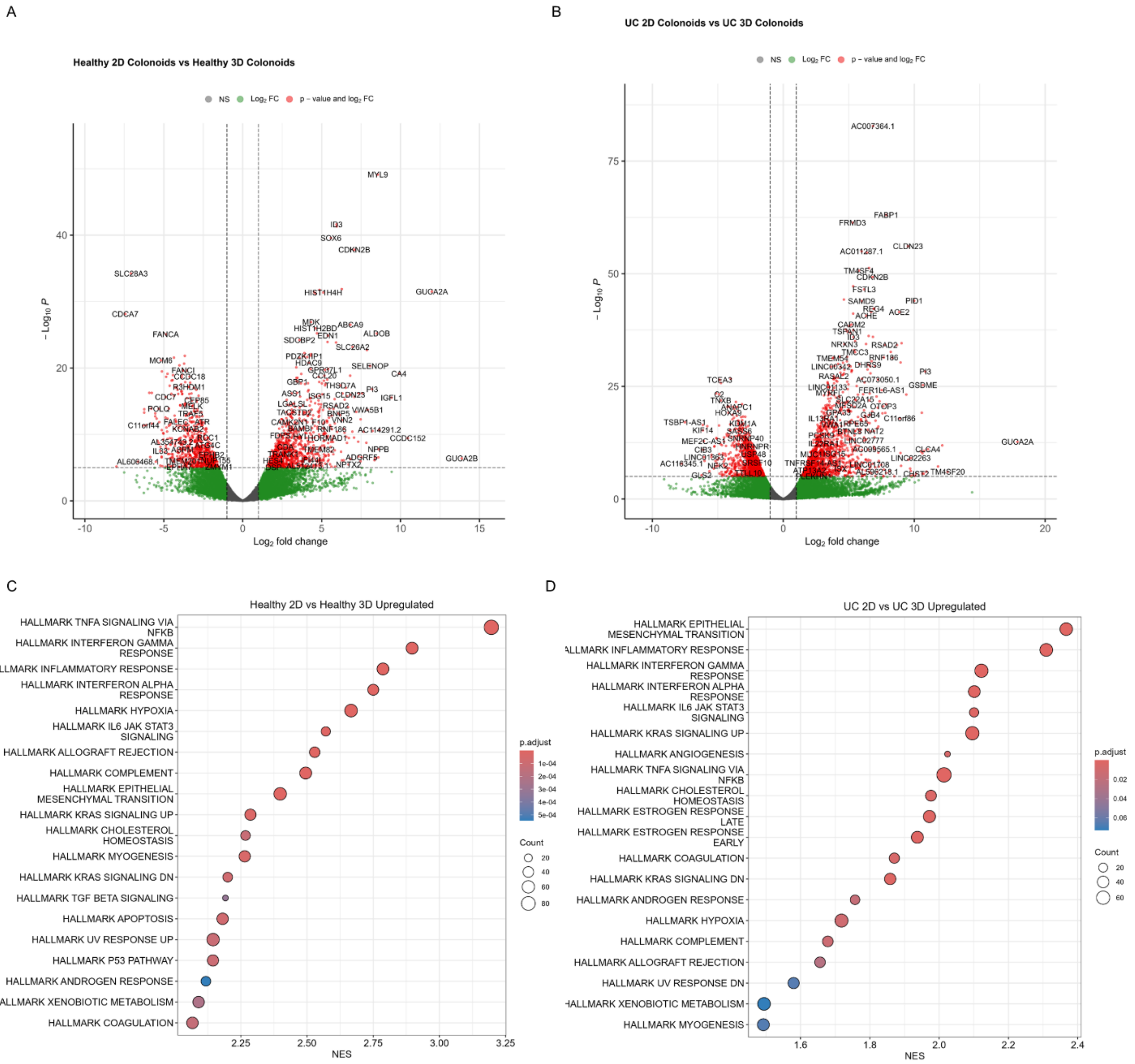


**Supplementary Figure 5.** Volcano plots for differentially expressed genes comparing 2D vs 3D colonoids from healthy (A) or UC (B) samples. Gene set enrichment analysis highlighting upregulated pathways on 2D vs 3D colonoids from healthy (C) or UC (D) samples.

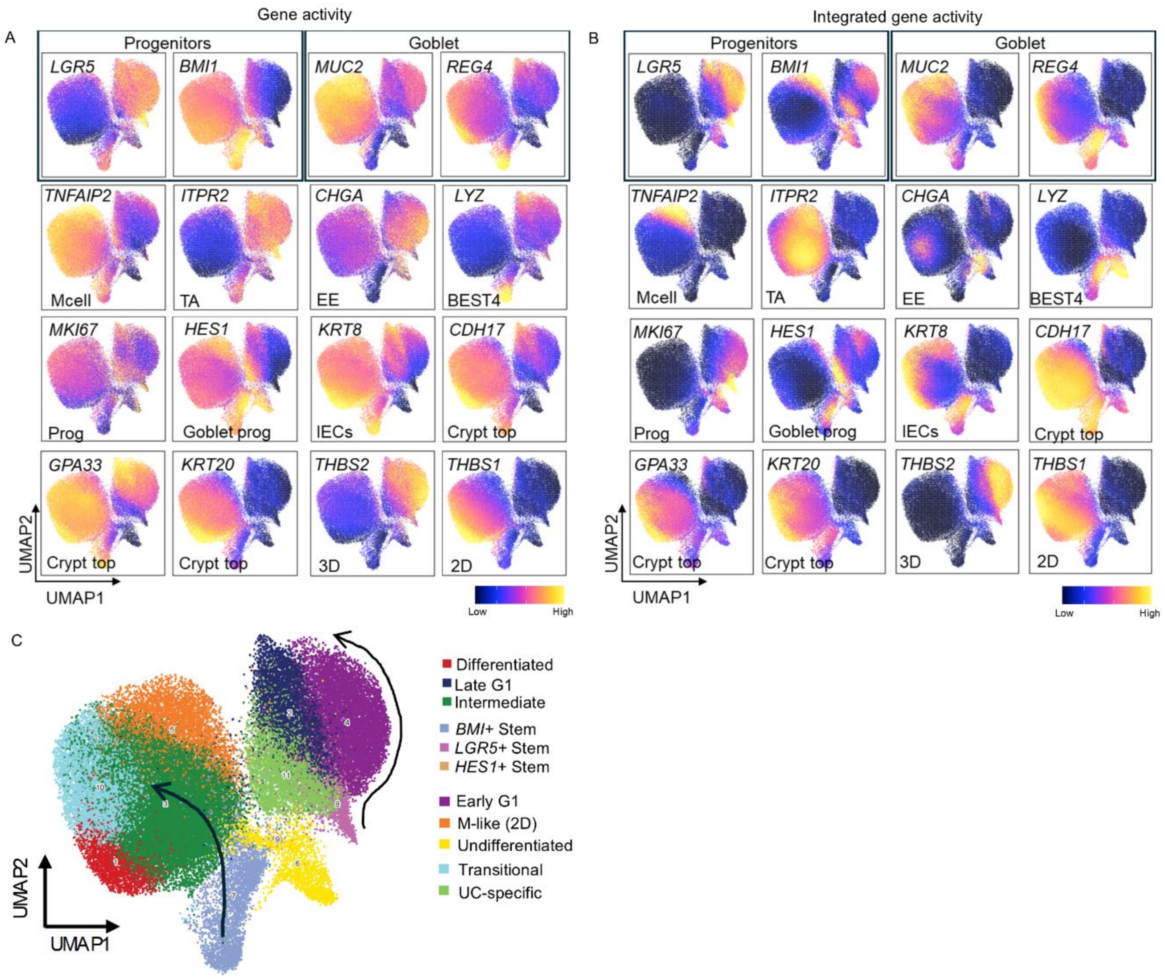


**Supplementary Figure 6.** A) RNA gene activity profiles of marker genes across intestinal cell types and colonoids. B) Integrated ATAC/RNA gene activity of marker genes across intestinal cell types and colonoids. C) UMAP plot of integrated ATAC/RNA-seq across all 3D and 2D colonoid samples.

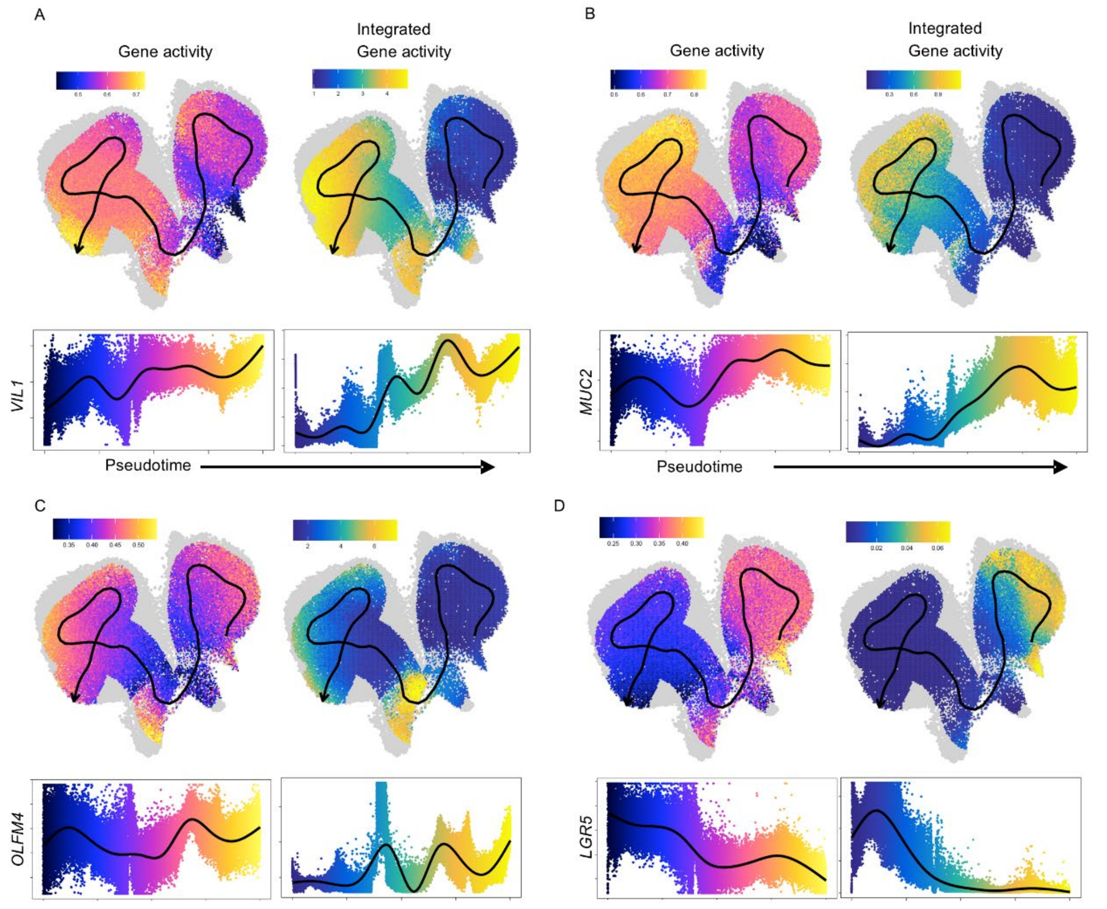


**Supplementary Figure 7.** Pseudotime-aligned expression dynamics of differentiation markers (A) *VIL1* and (B) *MUC2* and stem markers (C) *OLFM4* and (D) *LGR5* across all 3D and 2D colonoid samples.

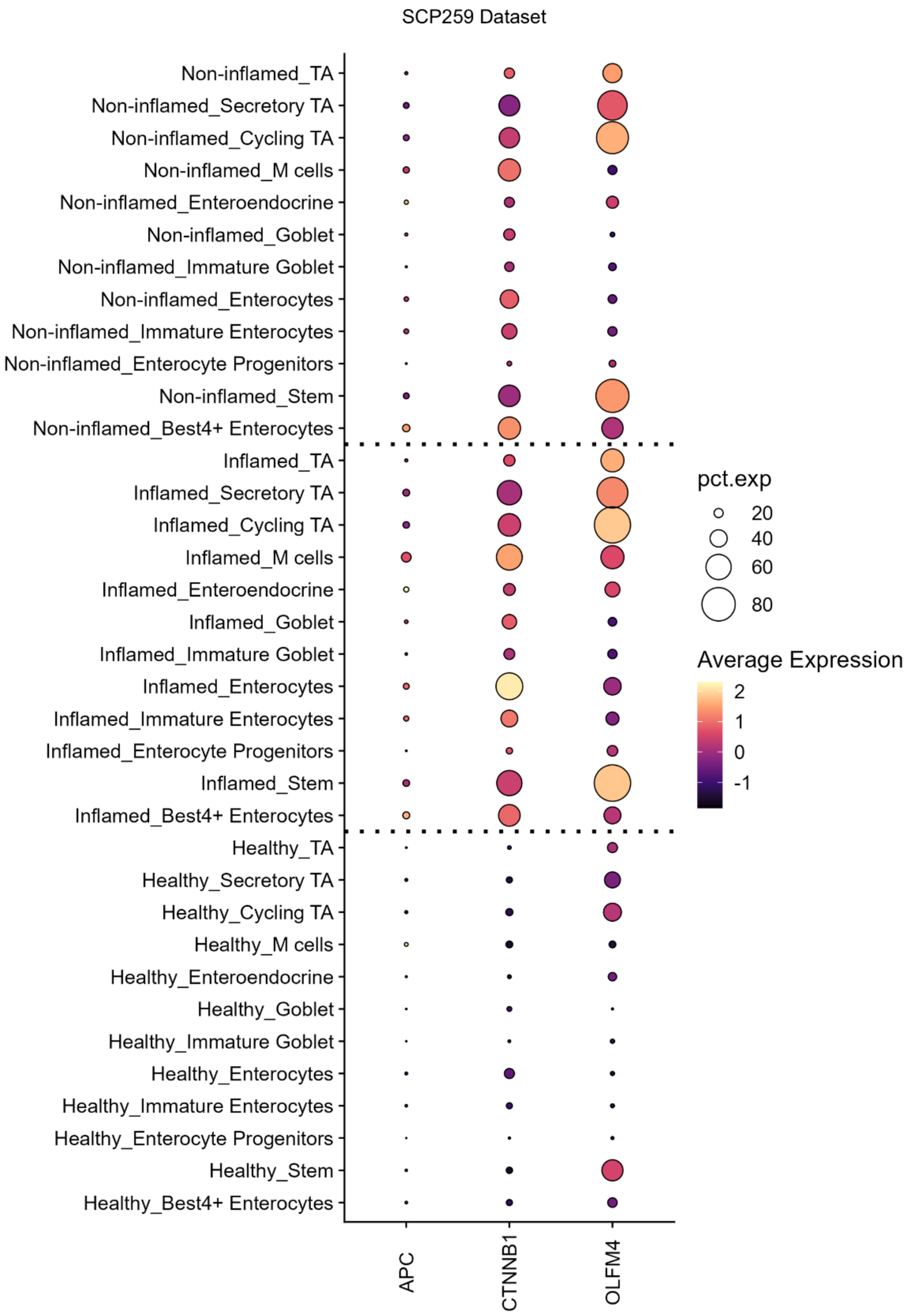


**Supplementary Figure 8.** Dot plot showing RNA expression of β-catenin (*CTNNB1*), *APC* ¸ and stem cell marker *OLFM4* in a public IBD scRNA-seq dataset (SCP259).

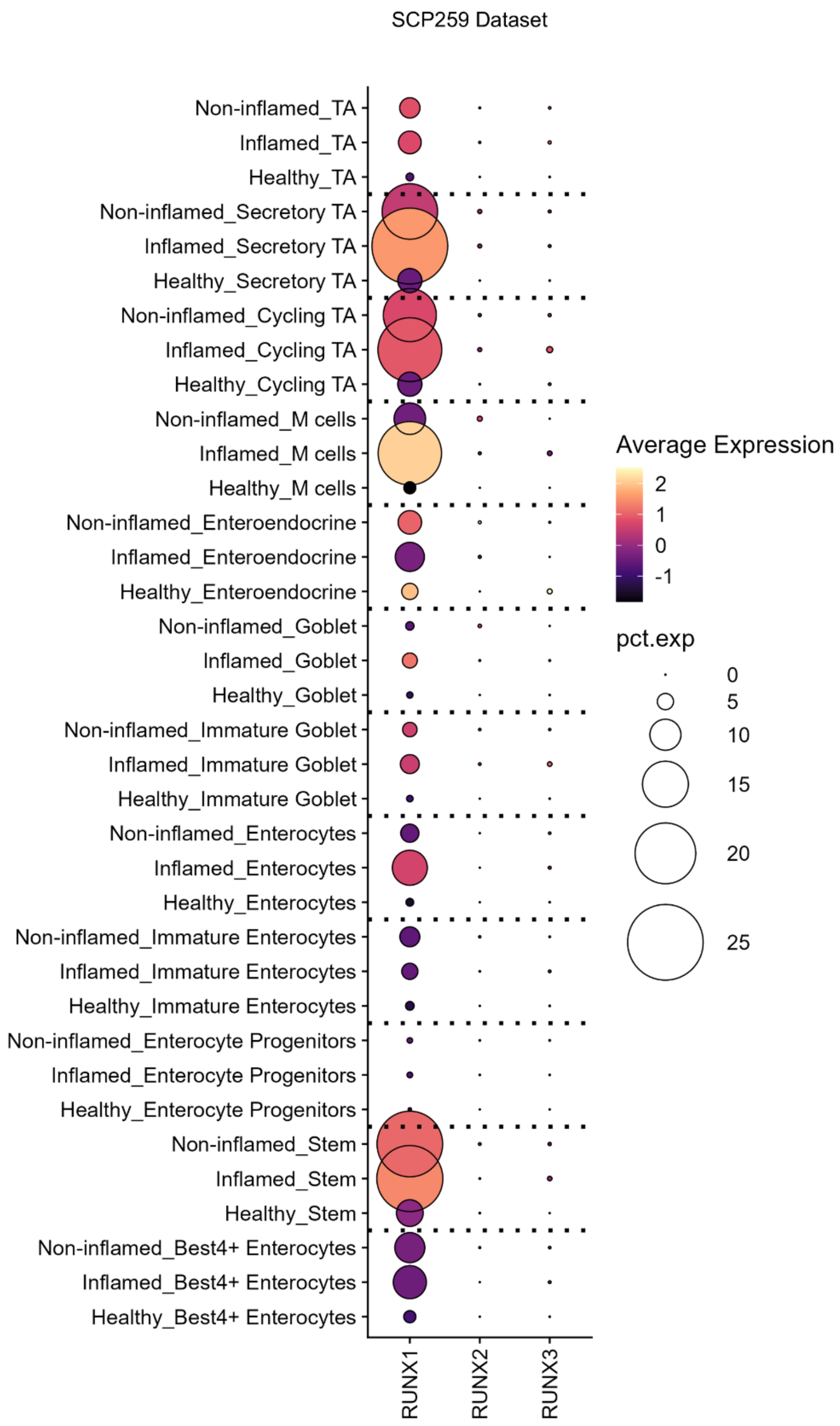


**Supplementary Figure 9.** Dot plot showing the expression of RUNX family members in a public IBD scRNA-seq dataset (SCP259).

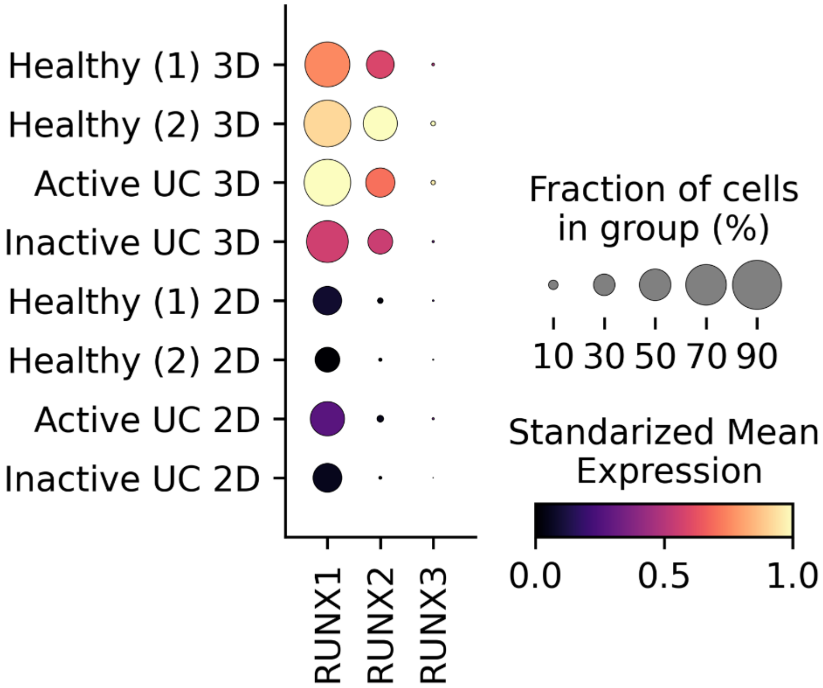


**Supplementary Figure 10.** Dot plot showing RNA expression of RUNX family members across all 3D and 2D colonoid samples.
